# Density-based longitudinal neuron tracking in high-density electrophysiological recordings

**DOI:** 10.64898/2025.12.19.695632

**Authors:** Yue Huang, Hanbo Wang, Jiaming Cao, Yu Chen, Xuanning Wang, Yujie Zhao, Hengkun Ren, Qiang Zheng, Jianing Yu

## Abstract

Tracking single neurons across days in high-density extracellular recordings is essential for establishing neural mechanisms of learning, memory, and post-injury recovery. However, in weeks-long recordings, identifying cross-day matches among thousands of units is confounded by changes in spike waveforms, unit turnover, and representational drift in neural responses. We introduce DANT (Density-based Across-day Neuron Tracking), an unsupervised framework that jointly estimates probe motion and neuron identity by alternating between density-based clustering in feature space and probe-motion correction inferred from provisional matches. Estimated drift is used to re-register spike waveforms across sessions, after which clustering is recomputed; this iterative loop continues until the set of matches stabilizes. In parallel, DANT learns a decision boundary from match and non-match assignments derived from the clustering results, enabling it to reject low-similarity candidate pairs. Applied to weeks-long Neuropixels recordings from the cortex and striatum in freely moving rats during stable behavior and task switching, DANT substantially increases match yield and reduces false negatives while maintaining a low false-positive rate relative to existing approaches. Together, these results indicate that DANT provides a general, unsupervised solution for longitudinal tracking in chronic, high-density recordings.

**The bigger picture:** Modern high-density probes such as Neuropixels record hundreds of units per session. When chronically implanted, many of these units can be stably recorded over extended periods, opening the door to tracking how large neural populations change over days to weeks, a question central to learning, memory, internal states, and recovery from brain injury. Yet, small shifts in probe position, changes in spike waveforms, and drifts in firing properties mean that linking units across sessions is challenging. DANT (Density-based Across-day Neuron Tracking) addresses this problem by providing an unsupervised framework that combines density-based clustering and iterative motion correction to identify cross-day matches with minimal subjective decisions. Applied to chronic Neuropixels recordings from freely moving, task-performing rats, DANT recovers longer and more complete chains of putative same neurons than existing approaches, while keeping the false-positive rate low. By making cross-day tracking more sensitive, scalable, and reproducible, DANT enables a wider range of neuroscience studies to examine the longitudinal behavior of single neurons and neural populations.

## Introduction

High-density electrode arrays such as Neuropixels probes can simultaneously capture the activity of hundreds to thousands of neurons. Chronically implanted Neuropixels probes often record neural activity for months^1–4^. Because of brain motion and interactions between external devices and the brain tissue^5^, some neurons are recorded across multiple days, whereas others become undetectable, and new units appear on the probe^3,6,7^. As noted previously^8^, this phenomenon presents “a problem and an opportunity.” Longitudinal single-neuron tracking enables studies of representational stability and how learning, experience, and injury shape neuronal representations in the brain^9^. On the other hand, within a single animal, recordings over multiple sessions with matched behavioral performance are often combined for group-level analysis, such as functional classification or encoding/decoding analysis^10^. Thus, it is necessary to reliably identify the same neurons across sessions and to avoid double-counting.

In the past, same-neuron matching has been performed with data from chronically implanted electrode arrays, for example, microwire arrays^11^, Utah arrays^8,12^, and tetrode arrays^13,14^. Tracking was typically based on examining similarities in features such as spike waveforms (1–4 channels), interspike intervals, firing rates, stimulus- or event-related spike modulations, and cross-correlations with other neurons. Pairs of units recorded between different sessions were identified as the same neuron when they showed highly similar features. Confidence in matches was established by comparing similarity distributions against null distributions (e.g., neuron pairs from different sites or animals)^8,14^. Among these features, waveform similarity is often the most critical, as all other features can naturally drift during learning and plasticity. Consequently, matching becomes more robust and reliable when waveforms from multiple closely spaced recording sites are available, as noted in tetrode recordings^14^, multi-site silicon probes^15^, and polymer arrays^16,17^.

Neuropixels probes have revolutionized neural recordings by oversampling spikes from single neurons across many closely spaced (∼20 μm) sites^3,4^. Furthermore, as brain movement is typically more pronounced along the probe’s axial axis than along the lateral axis, tracking waveform changes across multiple sites enables probe motion correction and subsequent waveform calibration using methods such as kriging interpolation.

Several methods leverage these high-density recordings to track neurons across sessions. One straightforward method concatenates pairs of sessions and sorts them jointly^3^. Sorting pairs of sessions together has the advantage of using the same spike-sorting standard (e.g., the same templates in Kilosort). However, the success of this method depends on the motion correction, which requires sufficiently similar activity patterns along the probes between two sessions. Practically, this approach also adds substantial burden to data processing, analysis, and storage, limiting scalability. Two independent alternatives have been proposed: the Earth Mover’s Distance (EMD) method that minimizes a combined cost of physical distance and waveform dissimilarity between matched neuron pairs^18^ and UnitMatch, which combines multiple features to train a Bayesian classifier using same-session recordings^7^. Both methods operate on pairwise comparisons across sessions.

Here, we introduce DANT, a clustering approach that leverages HDBSCAN (Hierarchical Density-Based Spatial Clustering of Applications with Noise)^19^. Briefly, we convert a weighted sum of similarities in features like spike waveforms and autocorrelograms into a pairwise distance that quantifies dissimilarity. Units recorded across sessions from the same neurons are presumed to have smaller distances in feature space and to form high-density clusters, identified by HDBSCAN. That is, this clustering-based approach extracts units from the same neurons across all sessions simultaneously. Probe motion correction and clustering are performed iteratively to enhance matching accuracy. We evaluated DANT using comprehensive datasets from freely moving rats performing reaction-time and self-timing tasks^20^, with chronic Neuropixels recordings from the striatum, motor cortex, and medial prefrontal cortex. Our results demonstrate that DANT enables rapid longitudinal tracking of the same neurons in freely moving behavioral paradigms with high fidelity over dozens of sessions simultaneously.

## Results

### Pipeline overview

The DANT pipeline is illustrated in Figure 1. Briefly, after spike sorting each session (1, 2, …, *n*) independently with Kilosort 2.5^3,21,22^, we assign each well-isolated unit within that session a unique ID (*k* = 1, 2, …, *K*; Module 1, m1; see Figure 1). Features for similarity analysis are extracted from each unit (m2), including spike waveforms across all channels, autocorrelograms (ACGs), and, optionally, peri-event or peri-stimulus time histograms (PETHs/PSTHs) that capture functional properties. Similarity scores between unit pairs are computed using feature-specific weights (equal weights of 1/3 per feature by default, m3), which are then transformed into pairwise distances representing dissimilarity. A density-based clustering algorithm applied to these distances identifies matched units hypothesized to originate from the same neuron (m4). Using these provisional clusters, linear discriminant analysis (LDA) optimizes the feature weights to maximize discrimination between matched and unmatched pairs (m5), yielding updated similarity scores and distances (m6). Clustering is iterated until the weights stabilize and the results converge. Using matched pairs’ spatial information, relative probe movement is inferred jointly across sessions (m7), and spike waveforms are remapped to probe recording sites, correcting for movement-induced changes in waveform distributions across channels (m8). After three iterations of m3– m8, the clustering output (m4) undergoes quality control (m9) to remove within-cluster pairs that fail the LDA-derived similarity criterion. Clusters are then assigned IDs (m10) representing units recorded across multiple sessions (from 2 up to *n*) that are hypothesized to originate from the same neuron.

**Figure 1.**
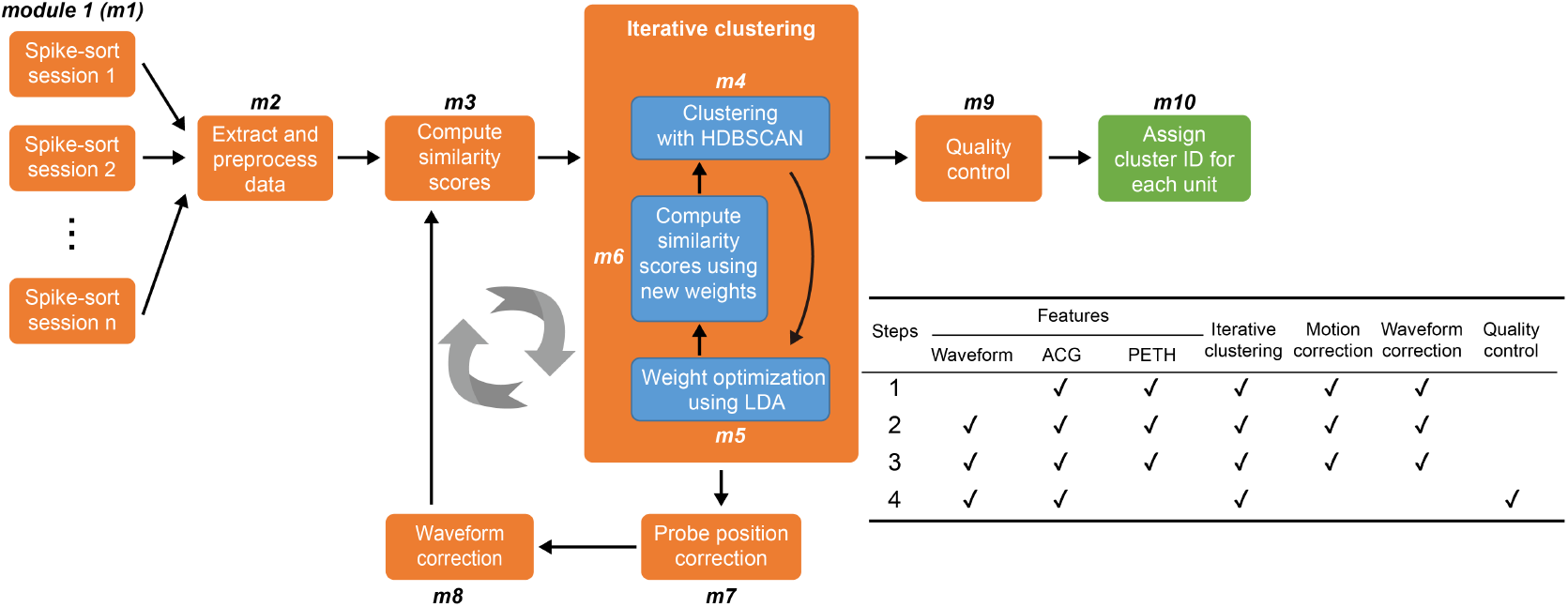
Outline of the DANT procedure. See the main text for details.

### Motion correction

In chronic preparations, even though the probe is rigidly fixed to the skull, displacements between the probe and neurons still occur due to brain motion and brain tissue responses to the implant^5,23,24^. These displacements reduce waveform similarity for the same neurons across sessions, complicating long-term tracking. In Neuropixels probe recordings, axial motion along the probe is captured by the probe’s densely packed channels. In the example shown in Figure 2A from a chronic recording in the secondary motor cortex (M2, see Figure 2C), the probe shifted upward relative to the recorded neuron (i.e., towards the brain surface) between day 1 and day m, resulting in a redistribution of waveforms across recording sites (Figure 2A1). Estimating this displacement (Δ*p*_1*m*_) allowed us to virtually restore the day-m probe position to its initial vertical location and to interpolate spike waveforms back onto the original recording sites (Figure 2A2). This correction increased waveform correlations between sessions from 0.89 to 0.97 (Figure 2B).

**Figure 2.**
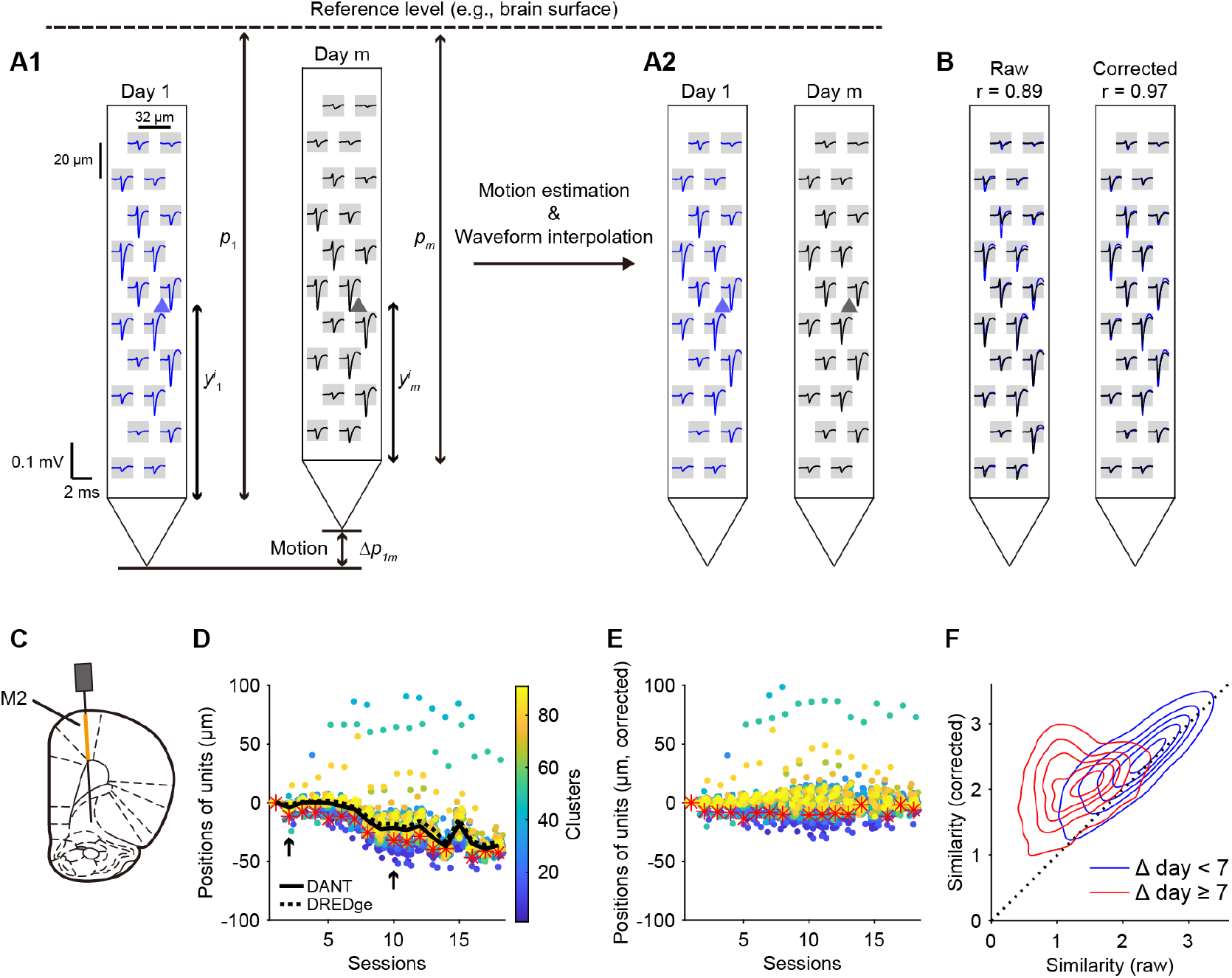
Probe motion correction. (A1) Schematic illustrating the relative displacement between the probe and the recorded neuron (triangles) across two sessions (days 1 and m), resulting in a redistribution of waveforms across recording sites. (A2) Illustration of waveform remapping following inference of relative probe motion. (B) Correlation coefficients of spike waveforms with and without motion correction. (C) Schematic of the recordings from the secondary motor cortex (M2) depicted in (A1–A2). (D) Relative positions of units on the probe. Unit positions in each session are normalized to their position in the first session in which they were detected. The red asterisk marks the unit shown in (A–B); arrows indicate two example sessions in A1. The thick black curve and dashed black curve denote probe position estimates by DANT and DREDge, respectively. Negative values indicate a decrease in unit positions relative to the probe tip, implying upward probe movement relative to the brain. (E) Motion correction minimizes the summed squared displacement of relative positions of all units. (F) Motion correction increases waveform similarity for matched unit pairs. Contours represent five density levels from 10% to 90% of the data.

The three-dimensional position of recorded units relative to the Neuropixels probe was estimated using a triangulation-based method (Equation 1)^25^. However, because the probe’s position across sessions was unknown, we treated it as a latent variable and estimated it iteratively. First, without motion correction, we obtained a provisional set of matched units across all sessions (Step 1 in Figure 1). Although this initial screening did not capture all matched units, it yielded a subset of highly similar units, forming the basis for probe motion estimation. Across sessions, matched units from the same neurons exhibited similar but not identical patterns of relative motion (Figure 2D; see also^18^; Figure S1 for other 12 datasets). We then applied an optimization procedure to derive probe motion values, minimizing the summed squared displacement of all matched unit pairs after correction (Equation 2, Figure 2E). Our estimated probe motion closely resembled that derived from an alternative method, DREDge^26^, which aligned spike amplitude distribution profiles along the probe’s channel sites to infer displacement (Figure S2A–D). With probe position corrected, spike waveforms were remapped (Equation 6, at the end of Step 1), similarity scores were recomputed, and clustering was performed again (Step 2), yielding a refined set of matched units that further improved probe position correction. After one additional correction step (Step 3), we performed a final clustering step (Step 4). Different feature sets were selected for each step (see the table in Figure 1), with details and rationale provided below. After motion correction, the waveform similarity between matched units increased overall (Figure 2F), facilitating classification of matched versus unmatched pairs using spike waveforms.

### Feature similarity and density-based clustering

For similarity analysis, we selected the following features and computed Pearson correlation coefficients, which were then Fisher *z*-transformed between each pair of units: (1) waveform similarity, **S**_WF_ (Figure 3A): correlation of waveforms across the channels (38 by default) closest to the unit; (2) autocorrelogram similarity, **S**_ACG_ (Figure 3B): correlation of spike-time autocorrelograms; (3) functional similarity, **S**_PETH_ (Figure 3C): correlation of peri-event or peri-stimulus time histograms (PETHs or PSTHs).

**Figure 3.**
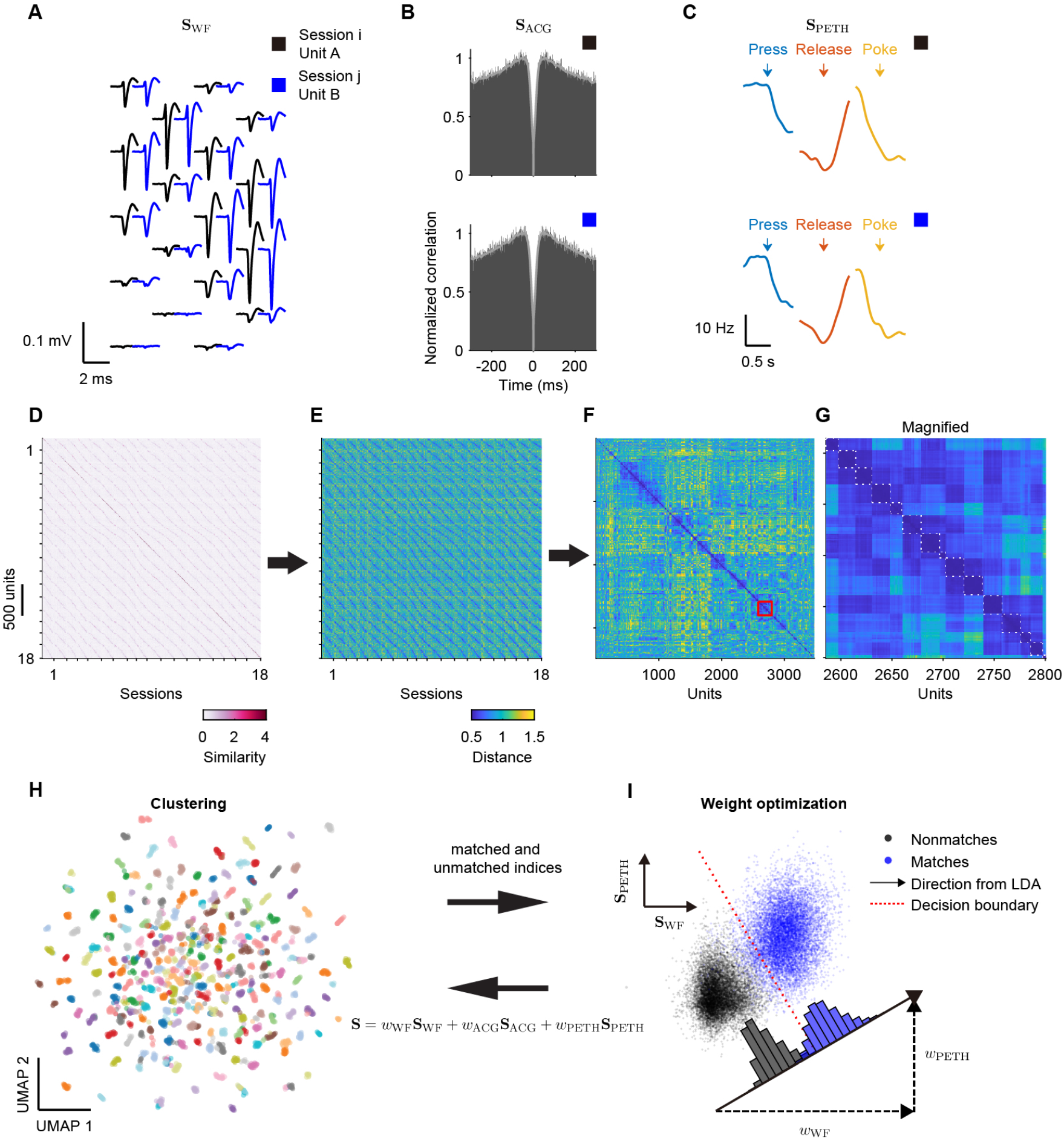
Density-based clustering. (A) Spike waveforms of two units from two different sessions. (B) Smoothed autocorrelograms of these two units, scaled to [0, 1]. (C) PETHs of two units, with neural activity aligned to behavioral events (lever press, lever release, and reward port nose-poke) in a reaction time task. (D) Similarity scores for all unit pairs, computed as the weighted sum of similarity scores from selected features. (E) Transformation of similarity scores into a distance matrix. (F) Density-based clustering via HDBSCAN identifies clusters of units with small distances (high similarity). (G) Zoomed-in view of the region marked by the red square in (F). Dashed squares highlight clusters identified by DANT that contain unit pairs with small pairwise distances (high similarity), indicating putative matches across sessions. (H) UMAP embedding visualizes unit clustering using pairwise distances derived from similarity scores; each point corresponds to a unit from one session. Colors indicate distinct clusters that DANT assigns to the same putative neuron across sessions. Only 20 colors are used, so the same color may appear in more than one cluster. (I) LDA separates matched (blue) from unmatched (black) unit pairs, optimizing feature weights to maximize discrimination. Dashed arrow lengths denote the LDA-derived weight for each feature (autocorrelogram similarity omitted for 2D visualization).

Using the Fisher transformation, we converted all correlation coefficients into similarity scores. All, or a subset, of these scores was then combined with weights (*w*_WF_, *w*_ACG_, *w*_PETH_) to form a single weighted similarity score (**S**, Figure 3D). Weight optimization was carried out iteratively together with clustering (Iterative clustering panel in Figure 1; Figure 3H,I).

A robust, unsupervised clustering method, Hierarchical Density-Based Spatial Clustering of Applications with Noise (HDBSCAN), was employed to group units that were highly similar in selected features^19,27,28^. This method has relatively few tunable parameters (notably the minimum cluster size and the minimum number of samples per core point) and makes no explicit assumptions about cluster shape, while accommodating clusters with different densities. To convert the similarity between units into a distance metric for clustering, the similarity score (**S**) was mapped to a tanh-transformed distance using (1+tanh(**S**)) ^−1^ (Figure 3E), such that highly similar pairs corresponded to small distances and dissimilar pairs to large distances in [0.5, ∞), which served as the input to HDBSCAN. HDBSCAN then examined the pairwise distance matrix and grouped units into clusters (Figure 3F), forming groups of units that were likely from the same neuron over sessions (Figure 3G, dashed boxes). Compared to EMD and UnitMatch, which perform pairwise comparisons of units and return matching results for one pair of units from two sessions at a time, the clustering approach yields tracking results for all units simultaneously across all sessions.

The quality of clustering is shown in a two-dimensional space using Uniform Manifold Approximation and Projection (UMAP), which approximately preserves pairwise distance relationships among units (Figure 3H). The clustering provides a binary—matched vs. unmatched—tag to each pair of units, allowing us to apply linear discriminant analysis (LDA) to find an optimal linear combination (weights) of features such that a one-dimensional boundary maximally separates these two groups (Figure 3I). The weights, which were initially set to equal values, were updated based on the results of LDA (Figure 3I). The similarity scores and pairwise distances were also updated, and the clustering was repeated. Clustering and weight updates were iterated until weights stabilized, which often occurred after just two iterations. Contributions to the weighted similarity scores varied across features. In Step 3 (Figure 1), the weights across 13 datasets were: **S**_WF_ = 0.51 ± 0.12, **S**_ACG_ = 0.13 ± 0.05, and **S**_PETH_ = 0.36 ± 0.11 (mean ± SD). In the final clustering step, the weights were: **S**_WF_ = 0.81 ± 0.07 and **S**_ACG_ = 0.19 ± 0.07. Excluding **S**_PETH_ avoids the influence of changing functional properties on tracking across sessions.

HDBSCAN is a hierarchical clustering algorithm that does not rely on a global density threshold for the data distribution. Instead, it groups data points into clusters based on their relative cohesiveness compared to surrounding points, irrespective of absolute distances or a global density^27^. Thus, points that are not truly from the same neuron can still be grouped into a cluster if they are relatively close to each other but distant from others, thereby producing false-positive matches. DANT further curates HDBSCAN outputs by imposing a similarity threshold defined by the LDA discriminant boundary (m9, Quality control). We identified two primary sources of errors. First, to avoid merging units from the same session, we enforced a constraint that each cluster contains at most one unit from any given session. Additional units from the same session exhibiting lower similarity scores were excluded (Figure S3A–E). Across 13 datasets, an average of 17.4 ± 14.1% of clusters (mean ± SD) contained at least one such excluded unit (2.4 ± 0.8 units per affected cluster, mean ± SD; Figure S3F,G). Second, for some clusters (16.3 ± 7.3%, mean ± SD), we applied a discriminant boundary derived from LDA to partition them into subclusters when subsets of units displayed sufficiently higher similarity scores among themselves compared to between subsets (Figure S3H–L). On average, these affected clusters were divided into 2.4 ± 0.2 subclusters (mean ± SD; Figure S3M, N).

### Stable and unstable functional representations in freely-moving, task-performing rats

We applied DANT to recordings from 11 rats performing lever-release simple reaction time (SRT) or self-timing (ST) tasks (Figure S4A). In the SRT paradigm, rats pressed and held a lever for a randomly selected foreperiod (FP: 0.75/1.5 s or 0.5/1/1.5 s), whereas in the ST paradigm, rats held the lever for at least as long as a fixed FP (0.75 or 1.5 s) and released it in the absence of a cue within a short time window (1 s). Although release initiation differed (tone-evoked in SRT versus self-initiated in ST), both tasks involved the same sequence of behavioral events—lever reach, press, release, and port nose-poke—providing a common structure for validating cross-day neuron matching based on functional properties. We recorded in consecutive sessions using Neuropixels probes, captured task events digitally, and synchronized high-speed videography (top and side views; side view shown in Figure S4B) via an infrared LED triggered by the stimulus.

For the example M2 recording (Figure 2C), the Neuropixels probe was implanted after the rat had learned the SRT task. After probe implantation, neural recordings were performed while the rat was performing the SRT task (sessions 1–5), during subsequent training on the ST task with two durations (sessions 6–14), and during retraining on the SRT task (sessions 15–18) (Figure S4E). Recordings spanned 18 sessions, during which neurons exhibited rich task-dependent modulation (Figure 4; Figure S4F). We tracked the rat’s forelimb position during lever-reaching (Figure S4C,D)^29^ and applied DANT to examine the functional stability of neurons recorded across sessions (Figure 4A–E). Units appeared and disappeared in different sessions (Figure 4A). We recorded a total of 3479 well-isolated units from 18 sessions (193.3 ± 23.3 units per session, mean ± SD) and identified 533 unique neurons. Out of these neurons, 56 (10.51%) neurons were recorded throughout all 18 sessions. The distribution of tracked length for each neuron is shown in Figure 4B.

**Figure 4.**
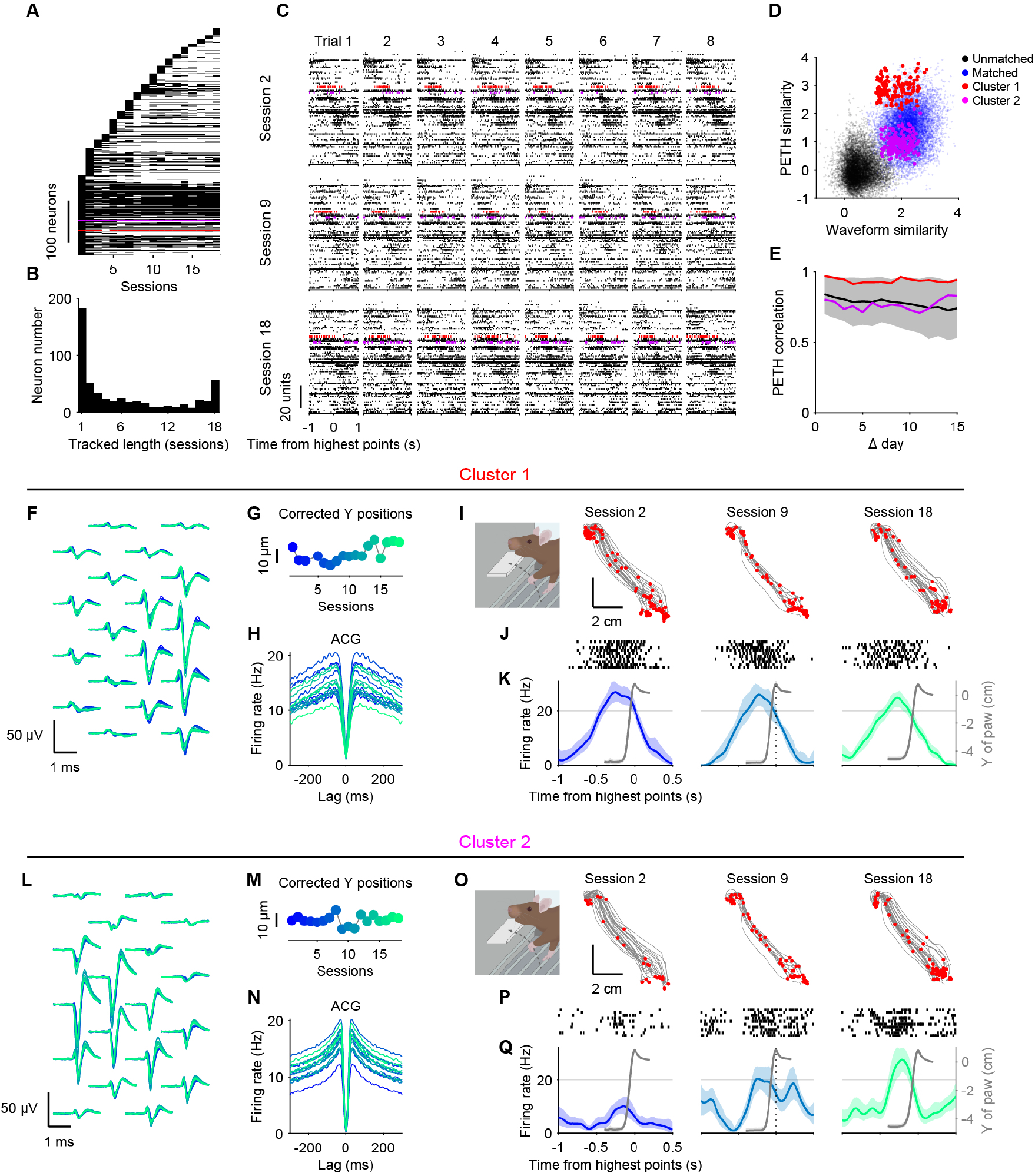
Example same-neuron groups in M2 recordings identified by DANT. (A) Presence of unique neurons across sessions, ordered by first appearance. Each row corresponds to a same-neuron group. Red and purple lines indicate two example neurons highlighted in subsequent panels. (B) Distribution of tracked duration (number of sessions) for all neurons. (C) Spike rasters of the same 99 neurons for eight consecutive example trials from each of three sessions. Only neurons tracked across these three sessions are shown, sorted by the time of their PETH peak in session 2. (D) PETH similarity versus waveform similarity for all pairs identified by DANT, with red and purple dots highlighting unit pairs from the two example neuron groups. (E) PETH correlation around the time of maximal paw height (lift-highest window: −1 to 0.5 s) plotted against the number of days separating the sessions. Data include neurons (*n* = 60) detected in at least 10 sessions with a mean firing rate above 5 Hz during the lift window. The black line is the grand average of all 60 neurons. The shaded area indicates the standard deviation of the average correlation for each day separation. Red and purple lines represent the correlation profiles for unit pairs within the two example same-neuron groups. (F) Overlaid waveforms after motion correction for an example neuron (Cluster 1 in (D). (G) Corrected unit position relative to the probe tip. (H) Smoothed autocorrelograms of spike times for the same neuron in individual sessions. (I) Left paw trajectories during lever reaches for 10 example trials across three sessions. The left panel shows a schematic of the rat lifting its paw to reach the lever. Red points mark spikes of the example neuron at the corresponding paw positions. (J) Spike rasters of the example neuron. (K) PETHs of the example neuron and vertical paw trajectories (gray), aligned to the time when the paw was at the highest point. Shaded areas indicate 95% confidence intervals of the firing rate. (L–Q) Same as (F–K), but for a different neuron (Cluster 2 in (D)).

When we focused on the lever-reaching stage—from 1 s before to 0.5 s after the peak of the reach trajectory—the spike patterns demonstrated overall functional stability in these neurons. The correlation coefficients of PETHs of matched units (60 unique neurons, 971 units in total) in this time window remained above 0.75 across more than 2 weeks of recordings. A stable neuron is shown in Figure 4F–K (Cluster 1); this neuron fired spikes before the initiation of forelimb lift and reached its maximal rate just before the forelimb movement onset (Figure 4K). Across different sessions (e.g., sessions 2, 9, and 18), PETHs aligned to the time of maximal reach were similar (correlation coefficient between sessions 2 and 18 = 0.950). Some units, however, showed changes in their functional properties over days. In a second example (Cluster 2, Figure 4L–Q), the coupling between spiking activity and forelimb lift was weak in session 2 but became stronger by session 18, yielding a PETH correlation of only 0.762 between those sessions.

Figure S4G–Q illustrates two additional neurons that fired during lever release from the same rat. Although response modulation depth exhibited modest session-to-session variability, the tracked neurons maintained consistent coupling to lever release, supporting the accuracy of our tracking method. Another recording example from the dorsolateral striatum (DLS) is shown in Figure S5 (Rat #8 in Figure S1). Neural activity was recorded while the rat performed the SRT task and during training on the ST task with two target durations (Figure S5A). Across 48 sessions, a total of 646 unique neurons were identified from 3404 recorded units (70.9 ± 14.6 units per session, mean ± SD; Figure S5B). Individual units were tracked for 24.5 ± 16.3 sessions (mean ± SD; Figure S5B,C). Figure S5E–I illustrates a neuron that increased its firing rate 0.5 s before lever press and peaked prior to the moment of lever press. Figure S5J–N shows another neuron that was activated in a short time window following nose-poke. Together, these data demonstrate the potential of combining DANT with chronic Neuropixels probe recordings to investigate how learning and experience shape neural dynamics in cortical and subcortical circuits during freely moving behavioral paradigms in rats.

We applied DANT to 13 datasets (four more example datasets in Figure S6A,B) and evaluated the persistence of tracked neurons across time using two metrics: (1) the matching probability and (2) the tracked length. The matching probability *P*_match_(*m*) quantifies the likelihood that the same neuron can still be identified across an *m*-day interval. For each pair of sessions separated by *m* days, we computed *P*_match_(*m*) as the fraction *n*_*i*+*m*_/*n*_*i*_, where *n*_*i*_ is the number of units recorded in session *i* and *n*_*i*+*m*_ is the number of units recorded in session *i*+*m* that were matched to units in session *i* (Figure S6C). Across datasets, the matching probability was 41.9 ± 15.9% (mean ± SD, *n* = 13 datasets) for *m*=7 (1 week) and 30.6 ± 17.3% for *m*=14 (2 weeks). The tracked length *t*_*i*_ of neuron *i* was defined as the total number of sessions—continuity not required—in which it was identified. For *n* recorded neurons with tracked lengths {*t*_1_, *t*_2_, …, *t*_*n*_}, we computed a survival function *S*(*k*) as 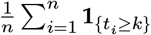, describing the probability that a randomly selected neuron appears in *k* or more sessions. Across datasets, 48.6 ± 24.9% of neurons were tracked for at least 7 sessions and 38.1 ± 18.4% of neurons were tracked for at least 14 sessions (Figure S6D).

To verify matches identified by DANT, we manually evaluated the matches in a subset of datasets by examining PETHs. Because neurons are expected to exhibit similar functional properties across neighboring days, we visualized and inspected the PETHs corresponding to each unit in every cluster identified by DANT using a custom Python graphical user interface (GUI, Figures S7, S8). We examined the results of datasets from motor cortex (*n* = 6) and DLS (*n* = 2), as neurons in these regions typically show clear tuning to one or a few task events and reasonable functional stability^13^ (Figure S4F).

In some cases, within a cluster, units showed markedly different functional properties, indicating errors in matching. For example, in Figure S7, 12 units recorded over 12 sessions were clustered by DANT (that is, 66 within-cluster “match” pairs). However, visual inspection revealed that these units in fact likely originated from two neurons with distinct functional preferences: the first 4 units fired during lever holding, whereas the remaining 8 units fired before the nose poke. The high similarity in spike waveforms and autocorrelograms was responsible for these units being mistakenly clustered as originating from the same neuron. Based on this manual assessment, there were 34 true-positive matches and 32 false-positive matches. For this cluster, the false-positive rate was therefore 32/66 (48.5%). We then split the cluster into two clusters consisting of 4 and 8 units, respectively. A majority of DANT-identified clusters, however, contained units with consistent functional properties, representing the same neurons recorded across sessions. For example, in Figure S8, all 57 units were correctly matched, and their functional properties—despite small drifts—remained similar across days.

Overall, most DANT-identified clusters were consistent with manual inspection. Across manually examined clusters (3113 clusters; Figure S9), the mean false-positive rate was 1.83 ± 10.43% (mean ± SD). Among all clusters, 2980 (95.7%) had no false-positive matches. These analyses highlight that false-positive matches still occur in DANT’s results, underscoring the value of post hoc inspection based on functional properties or other criteria to achieve high confidence that all units in a cluster originate from the same neuron. A further step in manual curation is to examine whether separate same-neuron groups, for example, different groups occupying consecutive blocks of sessions, can be merged according to well-defined criteria. This step reduces false-negative matches, which we analyze and compare with other methods in Figure 5.

**Figure 5.**
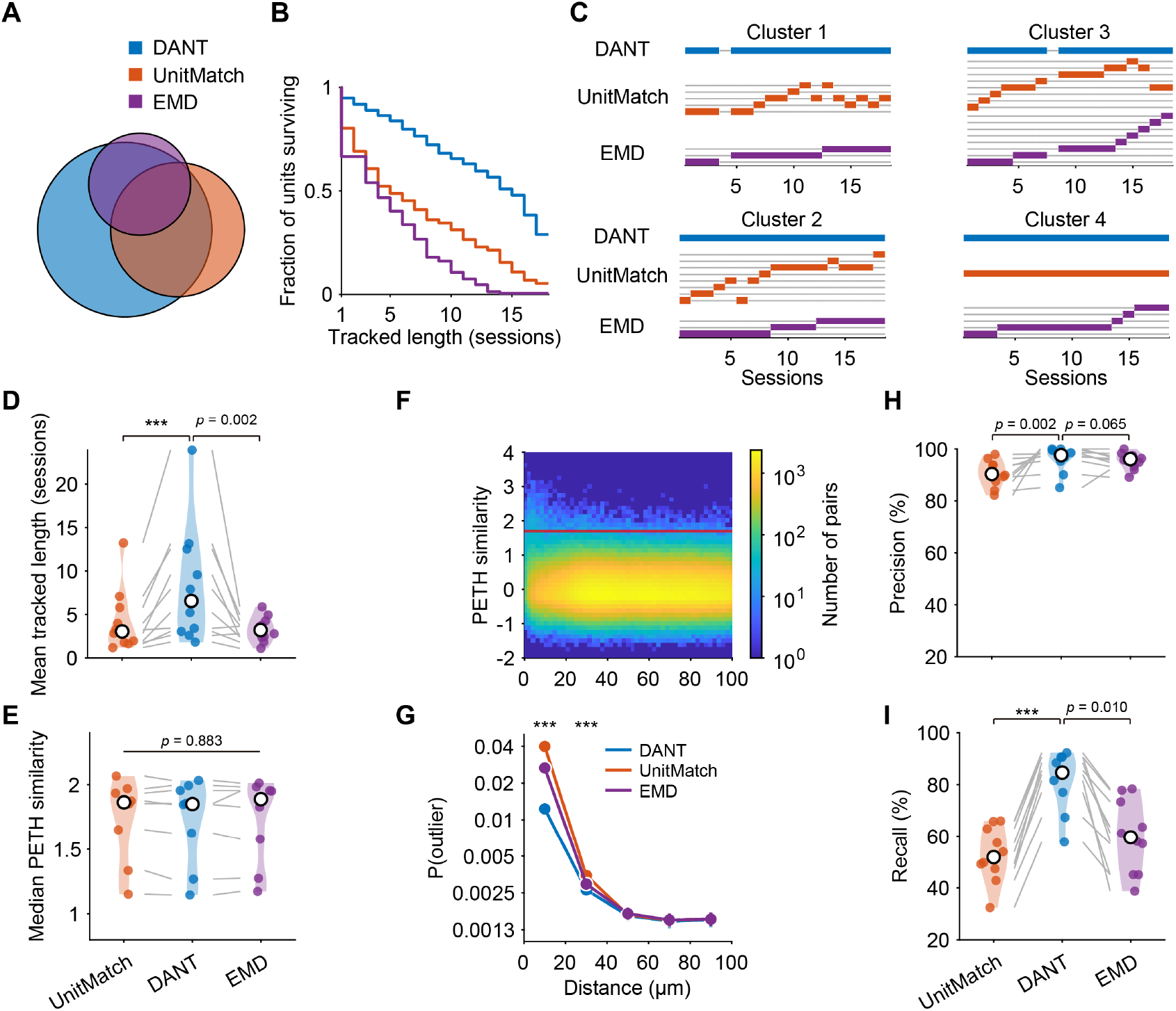
DANT increases matching yields compared to existing methods. (A) Venn diagram showing the overlap of matched units identified by the three different methods in an example rat. (B) Survival function showing, for each method, the proportion of units that could be tracked for at least *n* sessions in an example rat. (C) Tracking results for the example neurons shown in Figure 4F–Q (Clusters 1, 2) and Figure S4 (Clusters 3, 4) obtained with the three methods. Each gray line connects units that were grouped as the same neuron by a given method. (D) Mean tracked length for each method across 12 datasets. White points indicate median values. (E) Median PETH similarity between matched units recorded on consecutive days across 8 datasets. (F) Heatmap of PETH similarity for unmatched unit pairs identified by DANT across consecutive days, plotted as a function of inter-unit distance. Pixel intensity reflects pair counts. The red line represents the threshold used to identify outliers. (G) Proportion of outliers derived for each method, grouped into 20-μm distance bins. Error bars indicate 95% confidence intervals. (H, I) Precision and recall of each evaluated method compared to Kilosort-derived matches from 10 pairs of consecutive-day recordings. ***: *p* < 0.001

### Comparing to other methods

We compared DANT to existing methods. Currently, three approaches exist for tracking the same neurons with Neuropixels probes across sessions. One method performs spike sorting on pairs of consecutive sessions with motion correction at their junction^3^. This approach is computationally challenging for large numbers of sessions, requiring a dedicated spike sorting procedure for pairs of sessions in which units need to be compared. Additionally, it requires a continuous chain of matches for longitudinal tracking of a neuron across many sessions, constrained by the joint probability of matching a neuron across all consecutive sessions. If the chain is broken, the same neuron will be assigned to different matching groups and identified as different neurons. We thus focus on comparing our method to EMD^18^ and UnitMatch^7^, as these methods, like DANT, take results from spike sorting of individual sessions.

PETH features were excluded from the tracking procedures in all methods and used only to evaluate the tracking results. In an example dataset (an M2 recording), matching results derived from each method are shown in Figure 5A,B (the same dataset shown in Figures 2–4). DANT identified more matches and tracked units for longer periods (20057 matches, each neuron was tracked for 12.5 ± 5.7 sessions, mean ± SD) compared to UnitMatch (11848 matches, each neuron was tracked for 7.1 ± 5.6 sessions) and EMD (6857 matches, each neuron was tracked for 4.9 ± 3.9 sessions; Figure 5A,B). We observed that UnitMatch and EMD sometimes failed to track the same neurons across extended periods, instead fragmenting them into separate groups. For example, clusters identified by DANT in Figure 4F–Q and Figure S4H–Q were sometimes split into multiple groups by UnitMatch or EMD (Figure 5C).

We applied the three tracking methods to a total of 41591 units from 12 datasets across 10 rats. A linear mixed-effects model (Equation 19) was fitted to quantify neurons’ tracked lengths across methods. DANT yielded the longest tracked length compared with UnitMatch and EMD (DANT: 8.63 ± 1.80 sessions; UnitMatch: 4.50 ± 0.98 sessions; EMD: 3.63 ± 0.42 sessions, mean ± SE of estimated marginal means; Figure 5D). Post-hoc pairwise comparisons of estimated marginal means (Bonferroni-corrected) confirmed that DANT’s tracked lengths were significantly greater than both UnitMatch (*p* < 0.001) and EMD (*p* = 0.002). To assess the quality of these matches, we compared the PETH similarity between matched unit pairs detected by each method recorded on consecutive days. Our primary assumption is that matched units on consecutive days likely show high PETH similarity. Unit pairs with low firing rates and datasets with insufficient matched pairs (4 out of 12 datasets) were excluded to ensure stable median estimates with low variance. Under these constraints, matched pairs across all methods showed similarly high PETH similarity (*p* = 0.882, Friedman test; Figure 5E), indicating comparable quality of these matches.

In the absence of ground-truth data, examining PETH similarity between unmatched unit pairs helps compare false-negative rates across methods. In neighboring sessions, unmatched units either represent truly different neurons or the same neurons that a method failed to group, i.e., false negatives. If functional properties for a single neuron are relatively stable between two consecutive sessions, false negatives are expected to exhibit higher PETH similarity than pairs drawn from truly different neurons. A baseline distribution of PETH similarity for pairs that are extremely unlikely to be the same neuron was established by examining the PETH similarity between units that are widely separated in space along the probe^18^.

We examined the PETH similarity of unmatched unit pairs recorded across consecutive days and observed an increased proportion of pairs with high similarity at short inter-unit distances (Figure 5F), potentially reflecting false negatives. The PETH similarity for unit pairs separated by over 50 μm was modeled with a Gaussian distribution and a threshold, PETH_outlier_, was defined such that the probability of observing a value as large as or larger than PETH_outlier_ under this model was less than 0.01% (red line in Figure 5F) and such data points were treated as “outliers.” Our primary hypothesis is that a larger probability of detecting outliers when units were close to each other suggests a higher false-negative rate.

The proportions of outliers were then calculated in 20-μm distance bins for each method. Significant differences in outlier rates among methods were detected in the two closest bins (0-20 μm: *χ*^2^(2) = 1119, *p* < 0.001; 20-40 μm: *χ*^2^(2) = 19.92, *p* < 0.001; Figure 5G). Subsequent pairwise comparisons revealed that DANT yielded the smallest outlier rate (0.012) in the first bin compared to UnitMatch(0.04, odds ratio = 3.336, *p* < 0.001, Fisher’s exact test) or EMD (0.027, odds ratio = 2.195, *p* < 0.001). Together, these results indicate that DANT exhibits the lowest false-negative rate among all three methods, consistent with the examples illustrated in Figure 5C.

The proportions of outliers in DANT’s matching results can be further reduced by manual inspection. As described above, using a GUI, units can be excluded from a same-neuron group when they fail to meet predefined criteria, and different same-neuron groups covering non-overlapping sessions can also be merged when there is sufficient evidence to conclude that they originate from the same neuron, for example, when they show close proximity and similar PETHs. In four rats, manually merging separate same-neuron groups further reduced the outlier rate by more than half (from 1.58% among 9797 unmatched unit pairs before curation to 0.71% among 9632 unmatched unit pairs after curation) in the smallest distance bin (distance < 20 μm; Figure S9C–E), suggesting a corresponding reduction in the false-negative rate.

To further compare these three methods against a reference close to ground truth, we selected 10 consecutive session pairs from 5 rats (2 session pairs per rat) with high unit yields (two datasets per rat; 267.1 ± 121.7 units per session, mean ± SD) and jointly spike-sorted each pair with Kilosort 2.5^3^. These Kilosort-derived matches were used as a reference standard. We then compared the Kilosort-based matching with EMD, UnitMatch, and DANT. Note that EMD, UnitMatch, and DANT produced tracking results using all available sessions (*n*≥18 per dataset), whereas the Kilosort reference was obtained from paired sessions only. Precision (for each method, the proportion of its detected matches that were also detected by Kilosort; Equation 17) and recall (the proportion of Kilosort matches that were detected by each method; Equation 18) were calculated to quantify each method’s correspondence with Kilosort’s results. DANT achieved the highest precision (96.0% ± 4.8%, mean ± SD; *p* = 0.002 vs. UnitMatch, and *p* = 0.065 vs. EMD; Figure 5H) and recall (81.5% ± 11.2%, mean ± SD; *p* < 0.001 vs. UnitMatch, and *p* = 0.010 vs. EMD; Figure 5I).

To compare false-positive rates across different matching methods, we took advantage of a multishank Neuropixels 2.0 probe and the fact that units recorded on different shanks cannot originate from the same neuron, using a medial prefrontal cortex (mPFC) recording from Rat #3 in Figure S1 (sessions 4–10). We performed stable recordings over 7 days from two adjacent shanks (shanks 2 and 3) positioned at the same cortical depth. Each shank contained 192 channels spanning 1.425 mm. These recordings were characterized by high unit yields (162.1 ± 9.3 units on shank 2; 197.1 ± 12.8 units on shank 3; mean ± SD, *n* = 7 days) and minimal probe drift (maximum displacement of 35 μm, estimated by DANT).

We then generated a synthetic dataset by treating recordings from the two shanks as if they were recorded from the same shank on different “days” (Figure 6A). The x-coordinates of corresponding channels on shanks 2 and 3 were aligned to ensure identical channel maps in the synthetic dataset. Because units from different shanks are guaranteed to arise from distinct neurons, any matches between odd and even days in this synthetic dataset are, by construction, false positives and therefore provide an estimate of the false-positive rate. Because the EMD method requires neurons to be continuously present across sessions and was therefore not applicable, we only compared DANT and UnitMatch on this synthetic dataset. Both methods identified a large number of matched unit pairs (4926 matches, corresponding to 4.9 ± 2.2 sessions per tracked neuron on average for DANT; 3162 matches, corresponding to 3.5 ± 2.3 sessions per tracked neuron on average for UnitMatch; Figure 6B). Critically, despite DANT finding more total matches, it maintained a lower rate of cross-shank matches (3.82% for DANT and 4.87% for UnitMatch; *χ*^2^(1) = 5.02, *p* = 0.025, chi-squared test; Figure 6C). These results demonstrate that DANT yields more total matches without a corresponding increase in the false-positive rate compared to UnitMatch.

**Figure 6.**
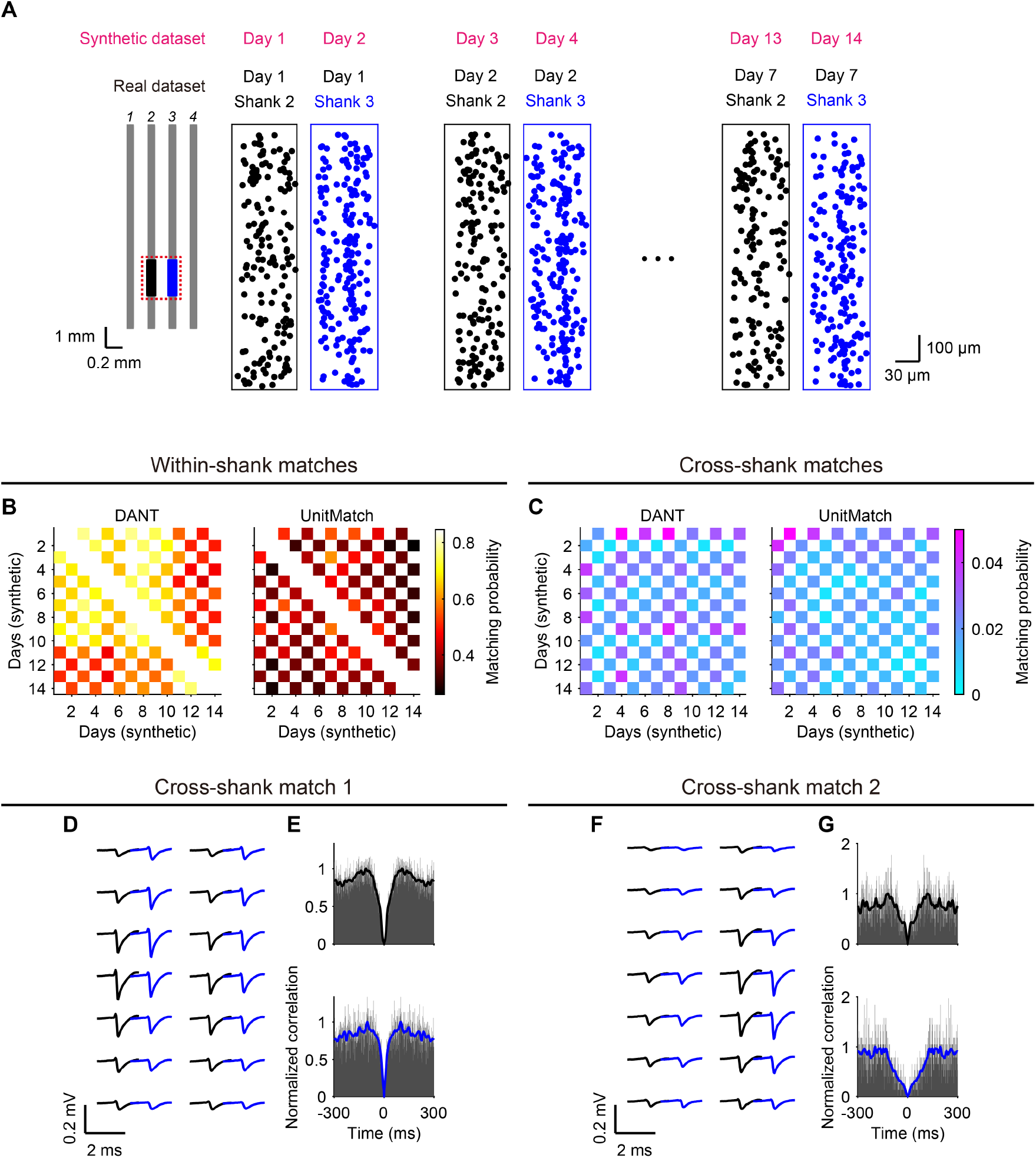
Performance of DANT and UnitMatch on a synthetic dataset. (A) Schematic of the synthetic dataset construction. Units recorded simultaneously from two different shanks (shank 2 and shank 3) in the same session were artificially treated as though they were recorded on separate “days”. By construction, matches between units from odd and even days therefore represent false positives. Black and blue points denote the locations of recorded units along shank 2 and shank 3, respectively. (B) Matching probability matrices showing within-shank tracking results of DANT and UnitMatch on the synthetic dataset. Pixel (i, j) represents the fraction of units from day i that were matched to day j. (C) Same as (B) for cross-shank tracking results. (D, E) Example of a cross-shank match identified by DANT. (D) Spike waveforms of the two units across corresponding channels. These units were recorded on shank 2 (black) and shank 3 (blue), respectively. (E) Normalized autocorrelograms of the two units. (F, G) Same as (D, E) for another cross-shank match.

Inspection of the cross-shank matches identified by DANT revealed highly similar waveform distributions across channels (Figure 6D–G). There is no reason to believe such highly-similar but false-positive matches will not occur for same-shank matches, which underscores the inherent difficulty of tracking the same neuron based solely on waveforms and spike-train patterns. For a set of *N* unit pairs that cannot correspond to the same neuron, assume the algorithm falsely matches any given pair with probability *r*, yielding *rN* false matches on average. In our synthetic dataset, all *n*_1_ cross-shank matches are false matches by definition. Among *n*_2_ same-shank matches, *rN*_2_ are expected to be false matches, resulting in a false-positive rate FPR_2_ =*rN*_2_/*n*_2_ for same-shank matches. The numbers of possible cross-shank and same-shank pairs, *N*_1_ and *N*_2_, scale with the number of session pairs; here, 7 ×7 = 49 cross-shank session pairs and 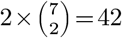 same-shank session pairs. We can then estimate the false-positive rate in same-shank matches, 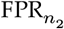, by the following approximation:

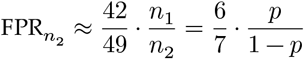

where *p* =*n*_1_/(*n*_1_ +*n*_2_) is the fraction of all matches that are cross-shank (3.8%). This yields a 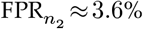, representing the false-positive rate expected for same-shank matches with DANT in this dataset.

This estimate, however, should not be generalized to other brain regions or recording configurations. The probability that two different neurons exhibit highly similar spike waveform distributions across dozens of channels depends on multiple factors, such as neuronal morphology relative to the probe and local cell density. In this particular analysis (Figure 6), the recordings were from mPFC, where the probe was approximately orthogonal to the principal axis of the dendritic arbors of pyramidal neurons.

Finally, DANT is the fastest algorithm compared to existing methods (Figure S10). This is likely attributed to the efficient implementation of its clustering method, enabling it to track over ten thousand units within an hour.

## Discussion

Large-scale neurophysiological recordings have expanded dramatically in recent years, particularly with the introduction of Neuropixels probes^3,4,30^, enabling simultaneous recording of hundreds of neurons. A central question is whether neurons recorded on one day retain the same functional properties, including their baseline firing rates, on subsequent days, and how these changes relate to the animal’s behavior or internal state^9^. These questions apply broadly across brain systems—including sensory, motor, emotional, and motivational circuits—and are of keen interest to many researchers. Accordingly, reliably tracking neurons longitudinally with chronically implanted high-density probes such as Neuropixels is highly desirable. Here, we present DANT, a fully unsupervised framework that uses density-based clustering as its central component and requires minimal subjective, manual decision-making. Using new datasets from chronic recordings in freely moving, task-performing rats, we show that DANT effectively clusters units that are likely to arise from the same neurons across days. Compared with existing approaches such as EMD and UnitMatch, DANT substantially increases match yield, while maintaining a false-positive rate comparable to UnitMatch (Figure 6C). In addition, an outlier-based proxy for false negatives, defined as the proportion of high-similarity unmatched pairs at short inter-unit distances, is reduced by approximately 50% with DANT compared with other methods (Figure 5G), suggesting that DANT likely produces fewer false-negative matches. Although DANT still produces some false-positive and false-negative matches (Figures S7, S9), our analyses indicate that, among the methods we evaluated, it provides a favorable trade-off between sensitivity and specificity and is also the fastest.

Unsupervised clustering has been a central tool for spike sorting based on examining waveform features^31^ and in some cases, the spatial location of spikes^32^. These features encode the similarity of spikes, allowing units originating from the same neurons to be automatically grouped using parametric algorithms, such as Gaussian mixture models^33^, or unsupervised clustering algorithms, such as distance-based methods like K-means^34,35^ or density-based methods^31,32,36–38^, including HDBSCAN^39^.

Tracking the same neurons over days in chronic recordings poses a similar problem. In some cases, spikes from multiple days are concatenated and spike-sorted jointly, so that units from the same neurons are naturally identified across sessions^3,15,40^. Alternatively, when each session is spike-sorted separately, units from different sessions must be compared post hoc using statistical tools. In previous work, similarity between two units across sessions is quantified by combining multiple feature-based similarity scores^8,12,14^. In some work^8,14^, these scores are then modeled as arising from a mixture of “same-neuron” and “different-neuron” Gaussian distributions, with the identity (same vs. different neuron) treated as a latent variable inferred by fitting a two-component Gaussian mixture using various EM-based algorithms. In other work, the same vs. different neuron identity for training data is predetermined^12^. In all cases, a classifier is trained on these predetermined or inferred identities and then applied to classify future unit pairs. In UnitMatch, which is designed for Neuropixels data^7^, a naïve Bayes classifier is initially trained on within-session data (similar to the approach in Dickey et al.^12^) and subsequently refined using putative matches after drift correction. It is then used to assign match probabilities to unit pairs across days.

DANT takes a conceptually different approach by using the unsupervised density-based clustering algorithm HDBSCAN to group all units from all sessions simultaneously. The clustering step yields matched vs. unmatched labels for all unit pairs; units within the same cluster are treated as matched, and units in different clusters as unmatched. We then train an LDA classifier on these labels to learn a decision boundary that separates matched from unmatched pairs and provides feature weights for combining multiple feature-based similarity scores tailored to each recording dataset. The resulting matches are fed into a probe motion correction step to improve waveform similarity estimation. HDBSCAN clustering, LDA training, and probe motion correction are iteratively combined to improve matching, and the final LDA decision boundary is used to curate the final clustering results. In this way, DANT extends the use of unsupervised clustering, which is popular in spike sorting, to the problem of tracking the same neurons over days.

Despite the robustness of the DANT framework for multi-day neuron tracking in Neuropixels data, it still produces both false-positive and false-negative matches. First, our analysis of a synthetic dataset (Figure 6) suggests that different neurons may have very similar waveform distributions and autocorrelograms. This type of ambiguity is inherent to extracellular recordings and, unfortunately, does not completely vanish even with dense channel layouts, contrary to earlier expectations^41^. In practice, putative same-neuron pairs can be curated post hoc based on additional criteria, such as whether the functional properties of units in a group remain reasonably stable between nearby sessions. In some systems, this assumption appears to hold^13,40^, whereas in other systems, the encoding properties of neurons are highly dynamic and can change rapidly with experience^42,43^. Thus, a general solution remains elusive: in such cases, examining the stability of functional properties is not a good criterion for identifying the same neurons. Overall, a dramatic change in neurons’ functional properties is either a genuine discovery or a sign of false-positive matches. Such decisions should be based on sufficient evidence and, where possible, control experiments. Second, because HDBSCAN prioritizes cluster “stability” rather than exact pairwise similarity, units with relatively low similarity may sometimes be grouped together, whereas units with high similarity may sometimes be separated. Third, probe motion of a long Neuropixels probe covering multiple brain regions including white matter can be complex, for example, non-rigid drift and displacement orthogonal to the probe can occur and distort waveform shape and distribution in a depth-dependent manner. To mitigate this issue, we incorporated non-rigid drift correction into DANT (Figure S11). Fourth, it is not uncommon to observe “disappearance-then-reappearance” of units in the same-neuron group (e.g., Cluster 5 in Figure S5). This pattern likely stems in part from the fluctuations in unit quality, whereby units sorted in some sessions may fail to meet the inclusion criteria for the DANT procedure. In many cases, these “missing” units can be recovered through manual inspection with a custom GUI and, when appropriate, integrated into the same-neuron chain.

Given these remaining limitations, researchers should critically assess match quality through manual inspection of DANT’s results, for example, using a custom GUI similar to that used for Figures S7 and S8. Manual inspection can help remove units that are incorrectly identified as matched based on specific criteria. Conversely, groups identified by DANT as different neurons may be merged if there is sufficient evidence that they arise from the same neuron, for example, if their spike waveforms are still similar and their functional properties are consistent. Ultimately, down-stream analyses should be tuned to the requirements of the specific scientific question. For example, when tracking the same neurons over time to study their dynamics, one may favor more conservative matching criteria to maximize confidence that all units are indeed from the same neuron; in this context, a short chain of highly confident same-neuron groups is often more useful than a long chain of less confident groups. By contrast, when combining unique neurons to build a pseudopopulation, one may instead seek to minimize the false-negative matching rate to maximize confidence that all included units are indeed from different neurons. Overall, DANT provides a practical and effective framework for tracking the same neurons across days, facilitating both longitudinal same-neuron analyses and pseudopopulation-based studies.

## Materials and Methods

### Animals

All animal procedures complied with animal care standards set forth by the US National Institutes of Health and were approved by the Institutional Animal Care and Use Committee of Peking University. Male adult Brown-Norway rats (*n* = 11), aged at least three months and weighing 250–400 g prior to water restriction and behavioral training, were obtained from Vita River (Beijing, China). Rats were housed individually or in groups of 2–3 on a 12-hour reverse light-dark cycle, with post-surgery housing limited to individual cages. During water restriction, water was earned through daily behavioral sessions of a simple reaction time or self-timing task (6 days a week); supplemental water was provided via manual delivery at least three hours post-session if intake fell below 12 mL. Rats’ body weights were monitored daily and maintained at approximately 80% of their free-feeding weight.

### Surgeries

The implantation of Neuropixels probes was performed in two stages, using 3D-printed recoverable implants designed by the Buzsáki lab^44^. Rats were anesthetized with 4% isoflurane for induction and maintained at 1.5–2% throughout the procedure, with vital signs monitored. In the first surgery, the dorsal skull surface was exposed and cleaned. Four to six skull screws were inserted to anchor the implant, and two ground screws were placed in the cranium above the cerebellum. Super-Bond C&B cement (Sun Medical Co., Ltd., Moriyama, Shiga, Japan) was applied over the skull, except at the implantation site, to attach the 3D-printed headcap. After at least two weeks of recovery, the second surgery was conducted.

In the second surgery, a craniotomy was performed, followed by probe implantation targeting the motor cortex, medial prefrontal cortex, and/or the dorsal striatum. The probe was glued to a recyclable drive (R2Drive, 3Dneuro)^44^ and lowered into the target areas at 1–2 μm per second using a manipulator. The craniotomy was sealed with Vaseline and bone wax to prevent leakage. Recordings began 3–7 days post-implantation, after full recovery and the stabilization of the probe. Unit activity was monitored in the homecage for 15 min daily from the second day after implantation to examine probe stability and unit yields. A custom-made cap, fitted to the headcap, secured the recording cable without a commutator. Recordings (*n* = 11 rats) were conducted daily for one-hour sessions over weeks in a simple reaction-time or self-timing task.

### Data processing

Data were acquired using SpikeGLX software (https://billkarsh.github.io/SpikeGLX/). For Neuropixels 1.0 probes, a 300 Hz hardware high-pass filter was enabled. Each session’s data underwent preprocessing via CatGT (https://billkarsh.github.io/SpikeGLX/help/catgt_tshift/catgt_tshift/), which applied phase-shift correction and global common average referencing. Preprocessed data were spike-sorted using a modified version of Kilosort 2.5 (https://github.com/jiumao2/Kilosort_2_5) that bypassed redundant preprocessing steps. Sorting results were manually curated using Phy (https://github.com/cortexlab/phy/) and custom plugins (https://github.com/jiumao2/PhyWaveformPlugin/) to exclude anomalous waveforms. In some large datasets, we applied quality metrics to automatically filter for well-isolated units using custom tools (https://github.com/jiumao2/AutoCurationKilosort). Following the definitions of quality metrics in Siegle et al.^45^, we included units only if they met all of the following criteria: (1) ISI violations < 0.05; (2) amplitude cutoff < 0.05; (3) presence ratio > 0.95; and (4) nearest-neighbor hit rate > 0.8. Units that failed any of these criteria were excluded from further analysis in DANT. Definitions of the ISI violations, amplitude cutoff, presence ratio, and nearest-neighbor hit rate metrics are provided in detail in^45^. Finally, raw waveforms for each accepted unit were reconstructed by “unwhitening” them using the whitening matrix generated by Kilosort 2.5.

Rats were initially trained in a simple reaction time task^20^, requiring them to press and hold a lever during a randomized foreperiod (0.75/1.5 s or 0.5/1/1.5 s) before releasing it in response to an auditory tone. Correct trials were rewarded with water accessed via port poking. Animals achieved expert-level performance prior to electrode implantation. Post-surgery, rats performed either the original task or a self-timing task variant where the foreperiod duration (0.75 or 1.5 s) remained fixed, requiring internal estimation of the interval and lever release within a 1-second response window. Lever-press, lever-release, and port-poke timestamps from correct trials were used to compute peri-event time histograms (PETHs; ±500 ms, 1-ms bins), smoothed with a Gaussian kernel (standard deviation: 50 ms). The position of the left paw when the rat reached for the lever was tracked with DeepLabCut^29^. About 300 frames were manually labeled and the DeepLabCut training was done with default parameters in the TensorFlow environment. For each video clip centered on lever press, we manually inspected all trials to verify tracking quality and excluded trials with occlusions or atypical movements (e.g., pressing the lever with the head).

For each unit, we compiled: (1) Mean waveforms across all recorded channels. (2) Spike times. (3) Session-specific metadata (date, behavioral context). (4) Probe channel maps. (5) PETHs for lever presses, releases, and port pokes. These data formed the input feature set for DANT’s tracking pipeline.

### Unit localization

The method for three-dimensional (3D) localization of units adapted a monopolar current source model^25^. Each unit’s 3D position is denoted as *x, y, z*. Channel positions *x*_*c*_, *y*_*c*_ on the probe plane follow SpikeGLX/Kilosort conventions, defining the position of a channel relative to the probe tip, with *z*_*c*_ = 0 (fixed at the probe’s 2D plane). Using a monopolar current source model, each unit’s position was inferred via

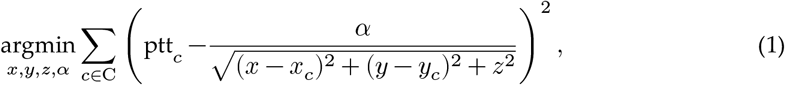

where ptt_*c*_ is the peak-to-trough amplitude of the spike waveform on channel *c*, C represents the indices of the *n* = 20 nearest channels to the peak channel (with maximum peak-to-trough amplitude), and *α* is the signal magnitude scaling factor. The optimization was performed using MATLAB’s “lsqcurvefit” function. The resulting position (*x, y, z*) denotes the relative position of each unit to the probe tip and was used to estimate probe motion, informing the motion correction algorithm described next.

### Probe motion estimation and correction

The method estimated probe motion across recording sessions using the location data of matched unit pairs, identified by the clustering algorithm described below. Let *N*_*s*_ be the total number of sessions, *N*_*p*_ the number of matched unit pairs, and *p*_*s*_ the scalar probe position (depth of the probe) in session *s* (*s*=1, …, *N*_*s*_; Figure 2A). For the *i*-th matched pair (*i*=1, …, *N*_*p*_), let 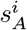 and 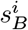 denote the sessions from which the two matched units were recorded, with 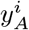 and 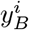 representing their spatial positions along the probe’s primary axis (relative to the probe tip, with 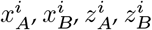 ignored; Figure 2A). The probe position vector across sessions **p** =(*p*_1_, …, *p*_*N*_) was estimated by solving:

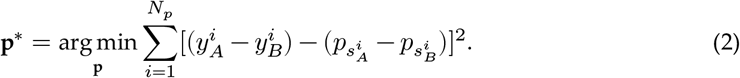

This optimization minimizes the mean squared relative displacement of matched units in the brain if the probe position is corrected using **p**^∗^. The optimization was performed using the “fminunc” function in MATLAB Optimization Toolbox. Estimated probe positions were then converted to relative values by subtracting either the first session’s position 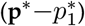 or the mean position across sessions 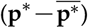, which were used for waveform correction and refined waveform correlation calculation.

DREDge^26^, a robust algorithm for registering extracellular electrophysiology recordings, was utilized to validate the probe motion. For the example rat shown in Figure 2, recordings from all sessions were concatenated and processed within the SpikeInterface framework^46^, which implements the DREDge algorithm. For the other example rats shown in Figure S2, which had longer recording sequences, only the first 10 min of each session was concatenated instead of the full recordings. Probe positions for each time point and spatial bin were estimated using the SpikeInterface “compute_motion” function with the “dredge” preset. The median of these estimated positions across all time and spatial bins was taken as the single, representative probe position for that session. To assess spike alignment, spikes detected by DREDge were binned in 5 s time intervals and 5 μm vertical spatial bins. The mean spike amplitude in each spatiotemporal bin was calculated. These values were plotted as a heat map (Figure S2B–D,F–H) to visualize the distribution of spike activity along the probe shank over time.

### Waveform correction

Let 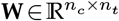 denote the matrix of the original average waveforms, where *n*_*c*_ is the number of channels and *n*_*t*_ is the number of time samples per waveform. Channel positions were represented in a two-dimensional coordinate system, with *x* denoting the lateral position and *y* denoting the depth along the probe shank. The original channel positions were collected as **v**_1_ = [*x*_1,*i*_, *y*_1,*i*_], *i*= 1, …, *n*_*c*_, where **x**_1_ and **y**_1_ are two vectors containing the *x*- and *y*-coordinates for each channel. Motion-corrected channel positions are defined as **v**_2_=[*x*_2,*i*_, *y*_2,*i*_], *i*= 1, …, *n*_*c*_. Here, the lateral positions remain unchanged (*x*_2,*i*_ =*x*_1,*i*_), while the depth coordinates are corrected as 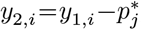, where 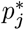 denotes the estimated vertical displacement in session *j*. Corrected waveforms, denoted 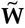, were estimated on these motion-corrected channel positions **v**_2_ using Kriging interpolation^47^. To quantify spatial proximity, distance matrices were defined as:

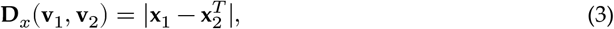

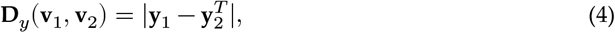

where **D**_*x*_(**v**_1_, **v**_2_) and **D**_*y*_(**v**_1_, **v**_2_) represent absolute differences in lateral (*x*) and depth (*y*) coordinates, respectively. Spatial similarity was modeled using an exponential kernel:

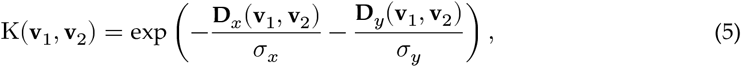

with spatial scale parameters *σ*_*x*_ = 20 and *σ*_*y*_ = 30 in μm. Finally, motion-corrected waveforms 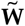 at channel positions **v**_2_ were obtained via:

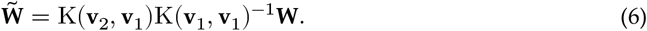

### Similarity scores

#### Waveforms

Waveform similarity between units was computed using raw or motion-corrected waveforms **W** from the *n* nearest channels to the channel with the largest amplitude for each unit. The similarity was quantified using Pearson’s correlation coefficient, followed by Fisher’s z-transformation. For units *i* and *j*, the *n* nearest channels to the largest-amplitude channel are denoted C_*i*_ and C_*j*_, respectively, based on their 2D spatial coordinates on the probe. The z-transformed correlations were computed as:

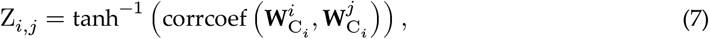

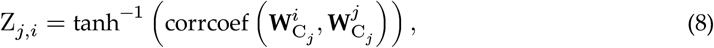

where 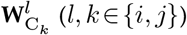 represents the waveforms of unit *l* on the channels indexed by C_*k*_. A symmetric similarity score was obtained by:

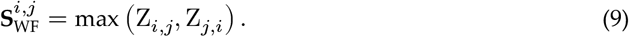

#### Autocorrelogram

The autocorrelogram for each unit was calculated with a maximum lag of 300 ms and a bin width of 1 ms. The resulting distribution was smoothed using a Gaussian kernel with a standard deviation of 5 ms and set to zero at lag 0. The autocorrelogram similarity score between units *i* and *j* was given by:

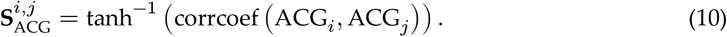

#### PETH

PETHs for each unit were precomputed during data processing, aligned to specific events in a forelimb-dependent simple reaction time task studied here. Three PETHs, corresponding to press, release, and poke events, were calculated with a time window of 500 ms around each event, using a bin width of 1 ms, yielding 1000 bins per PETH. The feature vector for unit *i*, denoted PETH_*i*_, was formed by concatenating these three PETHs into a vector of length 3000. The PETH similarity score between units *i* and *j* was given by:

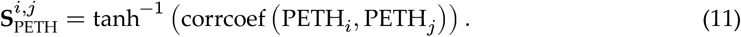

#### Weighted similarity score

The weighted similarity score between units *i* and *j* combined waveform (**S**_WF_), autocorrelogram (**S**_ACG_), and PETH (**S**_PETH_) similarity scores as a weighted average:

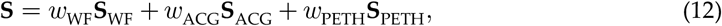

where weights *w*_WF_, *w*_ACG_, and *w*_PETH_ summed to 1, were initially equal (1/3), and were optimized for clustering accuracy. If PETH features were omitted due to task absence or significant response drift, the score simplified to:

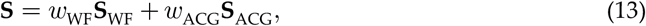

with *w*_WF_ +*w*_ACG_ = 1.

### HDBSCAN

Hierarchical Density-Based Spatial Clustering of Applications with Noise (HDBSCAN) is an unsupervised, hierarchical clustering algorithm based on the density of data^19,27^. This algorithm identifies groups of data based on local density profiles. To build a minimal spanning tree, HDBSCAN relies on the “mutual reachability distance”, defined as:

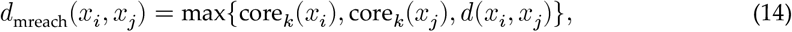

where core_*k*_(*x*_*i*_) is the distance to the *k*-th nearest neighbor of unit *x*_*i*_, and *d*(*x*_*i*_, *x*_*j*_) is a base distance (e.g., Euclidean). This metric ensures a density-dependent lower bound. To maximize sensitivity to small dense groups of units (including units from only two sessions), DANT sets *k* = 1.

In this study, the weighted similarity score **S** was transformed into a distance metric for HDBSCAN using the following heuristic formula:

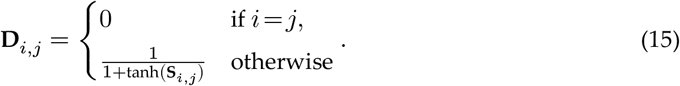

Here, **S**_*i,j*_ ∈ ℝ maps to tanh(**S**_*i,j*_)∈(−1, 1), yielding **D**_*i,j*_ ∈(0.5, ∞) for *i*≠*j*, with smaller distances approaching 0.5 for higher similarities. Chosen for its practical effectiveness in producing meaningful clusters in the context of chronic Neuropixels data, this heuristic transformation maps similarity scores to a distance metric suitable for HDBSCAN. The use of the hyperbolic tangent function ensures a smooth, bounded transformation, effectively capturing the relative differences in similarity for clustering units based on waveform or functional features.

The Python implementation of HDBSCAN^28,48^ was used, with parameters set to a minimum cluster size of 2 to allow small unit groups, a maximum cluster size equal to the total number of recording sessions, and minimum samples of 1 to maximize sensitivity to dense regions.

The resulting single-linkage tree was optimized using MATLAB’s “optimalleaforder” function to reorder units for visualization (Figure 3F,G). For further visualization, Uniform Manifold Approximation and Projection (UMAP) was applied to the precomputed distance matrix **D**_*i,j*_, producing a two-dimensional embedding that preserves local structure (Figure 3H). UMAP parameters were set to a minimum distance of 0.9, spread of 1, and 15 neighbors to optimize separation of clusters.

### Linear discriminant analysis (LDA)

The separation boundary between matched and unmatched unit pairs in the feature space was determined using linear discriminant analysis (LDA) with MATLAB’s “fitcdiscr” function. Only unit pairs within 100 μm along the probe’s y-axis were included in this analysis. We assumed that the feature vectors comprising waveform, autocorrelogram, and PETH similarity scores were drawn from multivariate Gaussian distributions with equal covariance matrices for the matched and unmatched classes. LDA yields a hyperplane **w**^⊤^**x**+*b*=0 that maximizes separation between these two classes. The discriminant vector **w** was normalized such that ∑_*i*_ *w*_*i*_=1, and the bias term *b* was scaled accordingly. The normalized coefficients of *w* were then used as weights to compute an optimized similarity score, integrating contributions from waveform, autocorrelogram, and PETH similarities (Equation 12). Similarity scores were projected onto this discriminant axis to distinguish matched from unmatched pairs. The scalar decision threshold, *s*_thres_ =−*b*, defined by the scaled hyperplane intercept, was subsequently applied during autocuration of DANT results.

### Iterative clustering algorithm

An iterative algorithm was developed to cluster Neuropixels unit pairs over *n*_iter_ iterations (10 iterations by default; Algorithm 1). A connectivity matrix **C** (*N*_*u*_ ×*N*_*u*_) was generated by HDBSCAN, with **C**_*i,j*_ = 1 indicating matched units. Similarity scores (Equation 12) were computed using default weights, transformed into a distance matrix, and clustered. HDBSCAN’s clustering results were used to identify matched and unmatched pair indices, which were then used by LDA to refine the weights. Updated weights were applied to recompute similarity scores for the next iteration, enhancing clustering accuracy.

For each dataset, three steps of iterative clustering (m4–m6), motion correction (m7–m8), and similarity score calculation (m3) were performed (Figure 1). Different feature subsets were selected for each step:

- Step 1: **S**_ACG_ +**S**_PETH_. This mitigates the impact of large probe drift between sessions, which could otherwise yield too few matched units.

#### Algorithm 1 Iterative HDBSCAN with Step-Specific Feature Selection and Autocuration

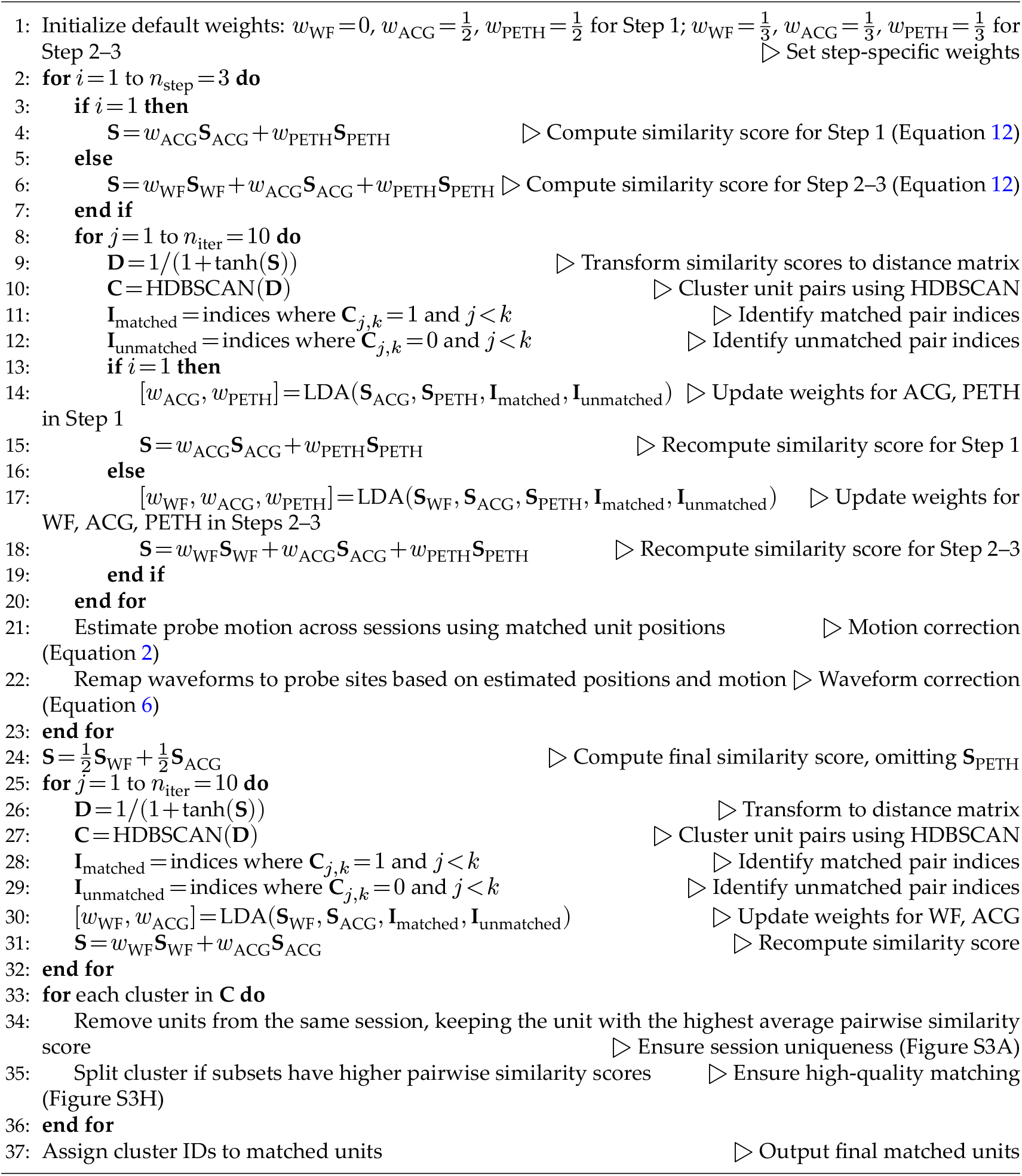

- Steps 2 and 3: **S**_ACG_ +**S**_PETH_ +**S**_WF_. This incorporates all features to optimize motion correction.

For final clustering, similarity scores were based only on **S**_ACG_ +**S**_WF_, omitting **S**_PETH_ to improve matching of units with significant functional drift.

Steps 1–3 were designed to infer probe motion and correct spike waveforms accordingly. The underlying assumption was that some units exhibited stable functional properties, reflected in **S**_PETH_ and **S**_ACG_, enabling robust motion correction in Step 1 to prevent large probe drift in a handful of sessions from disrupting clustering of continuously recorded neurons. Units with significant functional drift or minimal behavioral modulation contributed less to motion correction but were still likely matched in the final clustering step.

### Autocuration

Clustering results occasionally violated the constraint that units within a cluster must originate from distinct recording sessions. Units within each cluster were therefore inspected for session uniqueness, and if multiple units from the same session were detected, those with lower pairwise similarity scores to other units were removed.

Additionally, as the clustering algorithm prioritizes cluster stability over strict unit similarity, it can group units with low similarity. To ensure high-quality matching, pairwise similarity scores between all unit pairs within each cluster were evaluated, and the similarity threshold derived from LDA was applied. A connection was established between two units if their similarity score exceeded this threshold. Treating the resulting structure as a graph, its connected components were identified. If more than one component was detected within a cluster, it was split into subclusters corresponding to these components.

### Manual curation

To facilitate visualization and curation of clusters identified by DANT, we developed a Python-based program with a graphical user interface (available at https://github.com/jiumao2/DANT_UI). This tool extracts multiple features for each unit to enable systematic comparison, including motion-corrected waveforms, ACGs, PETHs, weighted similarity scores between units, and a custom figure illustrating the unit’s functional tuning within a given session (rasters and PETHs aligned to several events and separated by trial outcomes). The interface allows users to identify outlier units within clusters and reassign them to subclusters. In addition, it enables inspection of each cluster alongside its most similar clusters, based on the maximum weighted similarity score across units, to guide decisions on cluster merging.

Using this program, we manually curated clusters from eight datasets, comprising recordings from motor cortex (*n* = 6) and dorsolateral striatum (*n* = 2). Assuming functional properties remain consistent across neighboring days, units exhibiting clearly different functional characteristics were excluded from their original clusters.

### Neural activity stability across days

To assess the stability of lift-related neural activity across multiple days (Figure 4E), PETHs were computed for individual neurons. For the example rat presented in Figure 4, PETHs were generated for neural activity around the peak of paw lift (from −1 s to 0.5 s relative to peak lift). These PETHs were constructed with 10 ms bins and smoothed with a Gaussian kernel with a standard deviation of 50 ms. Only neurons reliably tracked across a minimum of 10 experimental sessions were included in this analysis (*n* = 152). To include only neurons significantly modulated around paw lifting, first, neurons with a mean firing rate below 5 Hz during the specified window were excluded. Second, a shuffling-based modulation test was performed for each neuron. Spikes were binned at 50 ms resolution around the lift-highest points (−1 s to +0.5 s) for each trial, and a PETH was computed by averaging spike counts across trials. The mean spike count across all bins and trials served as the baseline. Next, a random circular shuffling of spike counts was applied independently to each trial. A new PETH was computed from the shuffled data, and the summed squared difference from the baseline was calculated. This procedure was repeated 1000 times to generate a null distribution of modulation scores. The *p*-value for each session was defined as the fraction of shuffled scores that were greater than or equal to the observed score. To combine *p*-values across *n* sessions for each neuron, Fisher’s method was applied. Under the null hypothesis that PETHs from all *n* sessions are unmodulated (i.e., all *p*-values are random variables uniformly distributed on [0, 1]), the Fisher statistic −2 ∑ ln(*p*) follows a chi-squared distribution with 2×*n* degrees of freedom^49^. Clusters with combined *p*-values below 0.01 were included (60 out of 152).

For each included neuron, pairwise Pearson correlation coefficients were calculated between its PETHs from every possible pair of sessions, as a function of the temporal separation (in days) between those sessions. The mean and standard deviation of these correlation coefficients, aggregated across the remaining neurons (*n* = 60), were presented in Figure 4E.

### Time warping

Piecewise time warping was applied to align PETHs for lever-press, lever-release and port nosepoke on a common time axis (Figure S4F). For each trial *i*, let 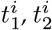 and 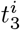 be the timestamps of the three behavioral events relative to lever press (so that 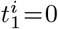). The median latencies of lever release and port poke relative to lever press are denoted as 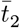 and 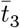, and 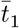 is set to 0.

Spikes were binned at 1 ms resolution, producing spike counts *x*_*i,j*_ at raw times *τ*_*i,j*_ in trial *i* and bin *j*. Within each segment (*k* = 1 for press to release, *k* = 2 for release to poke), any raw time 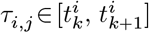 was linearly mapped to a warped time 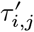 via

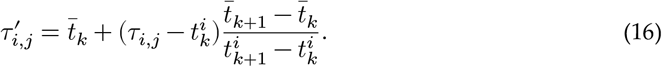

The warped spike pairs 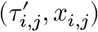 were linearly interpolated onto a common time grid using MATLAB’s “interp1” function. The time-warped PETH was then obtained by averaging the interpolated spike counts across trials and subsequently smoothing with a Gaussian kernel with a standard deviation of 50 ms.

### Comparison with UnitMatch and EMD

The example scripts provided in the GitHub repositories for UnitMatch (https://github.com/EnnyvanBeest/UnitMatch; MATLAB version) and EMD (https://github.com/janelia-TDHarrisLab/Yuan-Neuron_Tracking) were used for neuron tracking. Analyses were restricted to the same populations of well-isolated units across all methods. Default parameters for each method were applied across datasets without fine-tuning. For DANT, PETH features were excluded throughout the pipeline to ensure fair comparison. For UnitMatch, average waveforms for each unit were additionally extracted from the first and second halves of each session, as required by the algorithm. The “default” version of the UnitMatch algorithm was used to track the neurons over many recordings. UnitMatch failed in Rat #2 (16607 single units) and Rat #4 (14748 single units) due to an out-of-memory error, and these datasets were truncated to the first 31 sessions for successful processing. EMD encountered an error in Rat #6, and this dataset was excluded from further comparison. In total, 12 datasets were used for comparative analysis. All computations were performed on a computer with a 20-core Intel Xeon Silver 4114 CPU and 64 GB RAM.

To compare the PETH similarity of matched unit pairs recorded on consecutive days, similarity scores were calculated for matched pairs (Equation 11). Units with a mean firing rate below 1 Hz or a peak firing rate below 5 Hz in the PETH were excluded, since such pairs were more likely to yield artificially low similarity. Datasets with fewer than 100 valid matched unit pairs detected by either method were also excluded to ensure stable estimation of median similarity. The Friedman test was then applied to assess significant differences in PETH similarity across methods.

PETH similarity for unmatched unit pairs was evaluated as an indicator of potential false negatives. To establish a null distribution, we calculated PETH similarity for unit pairs recorded across consecutive days that were separated by more than 50 μm, which were assumed to represent correctly unmatched pairs. A Gaussian function was then fitted to this null distribution, yielding parameters μ = 0.025 and σ = 0.450. A one-sided threshold was then defined such that less than 0.01% of the null distribution exceeded it, corresponding to a similarity score of 1.70. Unit pairs with similarity above this threshold were classified as outliers. Subsequently, unmatched pairs were grouped into 20 μm distance bins. For each method, the proportion of outlier unmatched pairs within each bin was computed. Statistical comparisons of these proportions across methods were performed using chi-squared tests and when a signficant difference in distribution was found in a bin, pairwise Fisher’s exact tests were performed to compare different methods.

To further validate the tracking results, a subset of 10 high-quality, two-session recordings was selected from 5 rats. Data from these sessions were concatenated and jointly spike-sorted using Kilosort 2.5. For each unit identified in the individual session sortings, a corresponding unit in the combined dataset was sought. Units were considered “included” if more than 50% of their spike times (difference ≤ 1 ms) overlapped with a unit from the combined sorting. The majority of single units (87.9% ± 14.2%, mean ± SD) were successfully detected in this combined sorting output. For these identified overlapping units, those assigned to the same cluster by Kilosort were defined as a “match”. These Kilosort-derived matches served as the reference standard for validating matches found by DANT, UnitMatch, and EMD. Precision and recall were then used to quantify the agreement with Kilosort-derived matches. Precision was the proportion of detected matches that were also detected by Kilosort, whereas recall was the proportion of Kilosort matches that were detected by each method:

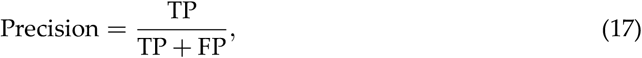

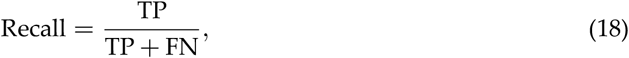

where TP is the number of matches identified by both Kilosort and the evaluated method; FP is the number of matches found exclusively by the evaluated method; and FN is the number of matches found exclusively by Kilosort.

To evaluate false-positive match rates, a synthetic dataset was constructed from a 7-day subset of data from Rat #3. These data were recorded using a Neuropixels 2.0 probe during a period of minimal probe motion. The same regions on two of the probe’s four shanks were consistently recorded across the seven-day experiment. These two shanks were 250 μm apart, thus minimizing the likelihood of recording from identical neurons. To simulate false-positive matches, units recorded from these two shanks on the same day were artificially assigned to separate “days” within the synthetic dataset, resulting in a 14-day dataset. DANT and UnitMatch were then applied to the synthetic dataset. False-positive matches were defined as pairs of units originating from different shanks.

### Statistics

All statistical analyses were performed in R. When comparing units’ tracked lengths across methods, a linear mixed-effects model was fitted using the “lme4” package with formula:

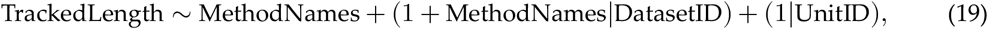

where DatasetID accounted for variability across datasets (including differences in session numbers and unit yields) by allowing both random intercepts and slopes, and UnitID accounted for repeated measures within units. Estimated marginal means were obtained using the “emmeans” package, and pairwise comparisons against DANT were performed with Bonferroni correction. For non-parametric comparisons of PETH similarity, precision, and recall across methods, the Friedman test was conducted using the “PMCMRplus” package followed by Nemenyi post-hoc pairwise analysis when appropriate.

## Acknowledgments

We thank Dr Jennifer Colonell for discussions and feedback on an early version of the manuscript.

## Funding

National Natural Science Foundation of China 32070983, 32271052, 32571183 (JY)

Natural Science Foundation of Beijing Municipality 5212007 (JY)

Peking-Tsinghua Center for Life Sciences (JY)

State Key Laboratory of Membrane Biology (JY)

## Author contributions

Conceptualization: YH, JY

Methodology: YH, JY

Investigation: YH, HW, JC, XW, YZ, HR, QZ, JY

Visualization: YH, JY

Supervision: JY

Writing—original draft: YH, JY

## Data and materials availability

Example data for the rat presented in Figure 2–4 are available via figshare (https://doi.org/10.6084/m9.figshare.30596258.v2)^50^ as part of the software demo. All additional data and code are available upon request.

## Code availability

DANT is available as both a MATLAB implementation on GitHub at https://github.com/jiumao2/DANT (GPL-3.0 license) and a Python implementation at https://github.com/jiumao2/pyDANT (GPL-3.0 license). It utilizes the HDBSCAN Python package for clustering^28^ (BSD-3-Clause license). Neuropixels data acquisition was performed using SpikeGLX at http://billkarsh.github.io/SpikeGLX/ (Janelia Research Campus Software Copyright 1.2). The Python application for manual curation of DANT results is available at https://github.com/jiumao2/DANT_UI (GPL-3.0 license). Spike sorting was conducted with a modified version of Kilosort2.5 at https://github.com/jiumao2/Kilosort_2_5 (GPL-2.0 license), followed by manual curation using Phy2 at https://github.com/cortex-lab/phy (BSD-3-Clause license) and custom plugins available at https://github.com/jiumao2/PhyWaveformPlugin (GPL-3.0 license). For a subset of datasets, automated curation was performed using AutoCurationKilosort at https://github.com/jiumao2/AutoCurationKilosort (GPL-3.0 license).

## Supplementary Materials

This PDF file includes: Figures S1 to S11

## Supplementary materials

**Figure S1.**
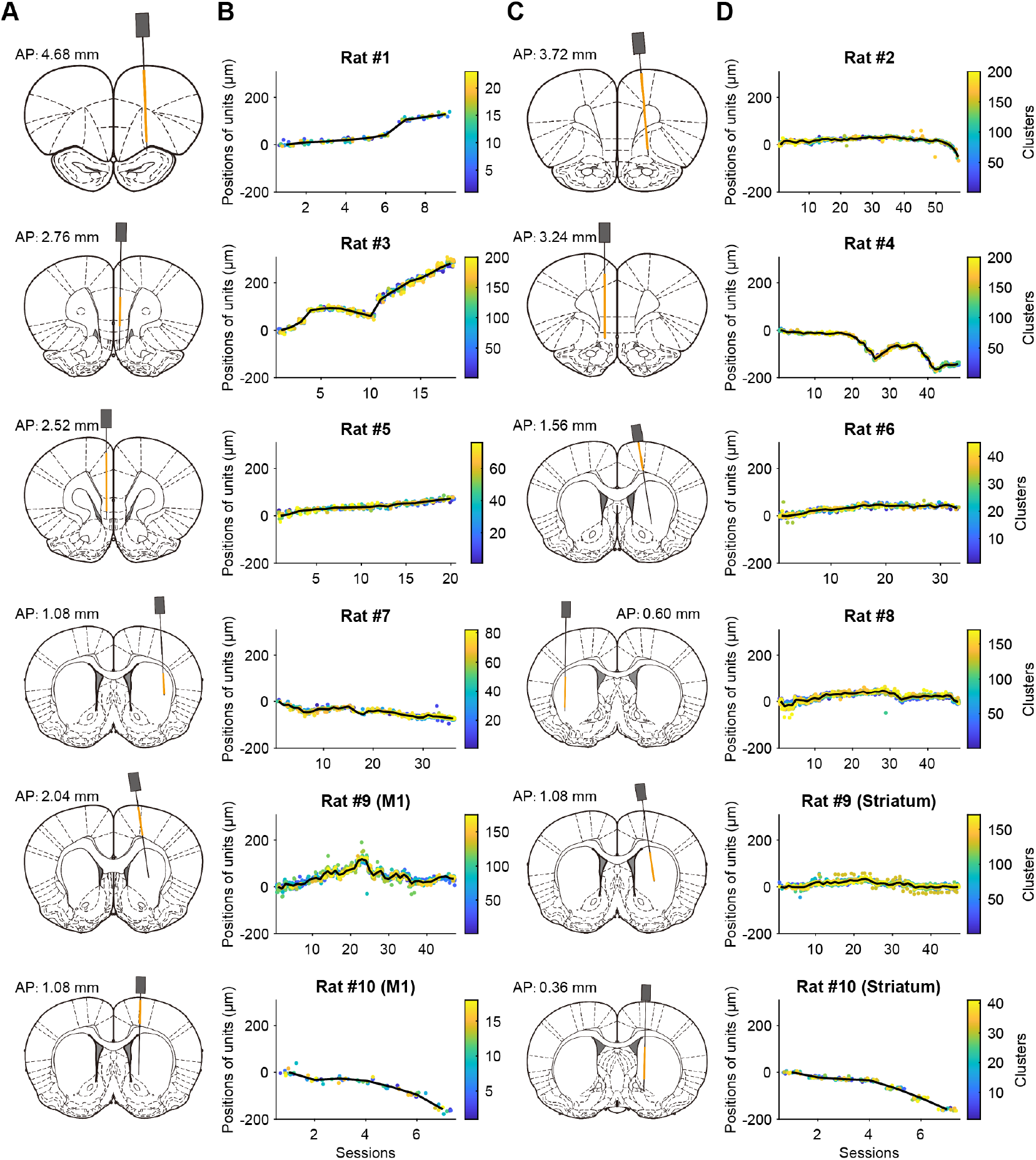
Probe motions across animals. (A, C) Schematics of the recording sites for each dataset. For multi-shank Neuropixels 2.0 probes, only one shank’s sites are shown and the other shanks targeted similar regions. Yellow shading denotes the span of recording sites included in the DANT analysis. In two rats (9 and 10), cortical and striatal recordings were analyzed as separate datasets since they exhibit different relative probe drifts. (B, D) Relative positions of units along the probe across sessions. The black line shows the estimated probe displacement relative to the first session. The positions of units in each detected cluster (up to 200 clusters shown) were normalized to the probe’s mean position across the sessions in which those units were recorded. Colors indicate cluster identity.

**Figure S2.**
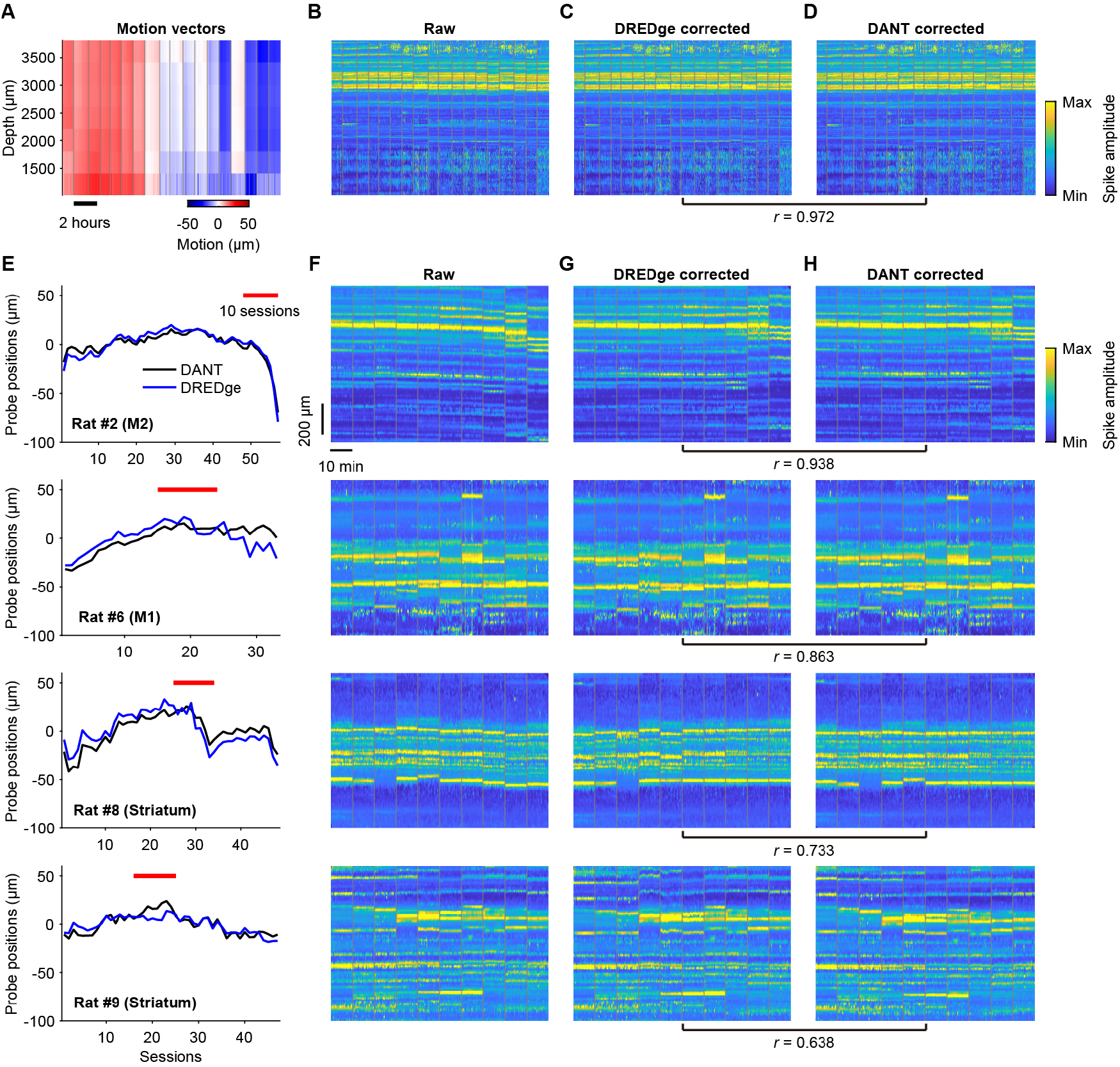
Comparison of DREDge and DANT for probe motion estimation. (A) Heatmap of probe motion estimated across depth and time by DREDge in an example recording spanning 18 sessions (same dataset as in Figure 2C–E), with color indicating displacement magnitude. Gray lines mark session boundaries. Probe position estimates from DREDge and DANT for this dataset are shown in Figure 2D. (B–D) Spike amplitude maps along the probe over time: before motion correction (B), after correction by DREDge (C), and after correction by DANT (D). Colors represent the mean spike amplitude within each spatiotemporal bin (min–max normalized across all sessions); gray lines mark session boundaries. Corrected spike amplitude maps are highly similar between C and D (the Pearson correlation, *r*, between the two maps is indicated below these panels). (E) Estimated probe positions from DANT (black) and DREDge (blue) in four additional example rats with recordings in cortex or striatum. For DREDge, median values of motion across depths within each session are shown. Red lines indicate the sessions displayed in (F–H). Probe motion is shown after normalization by the mean position across sessions. (F–H) Same as (B–D) for these four additional rats. Each session consists of a 10-minute recording segment used for motion correction with DREDge.

**Figure S3.**
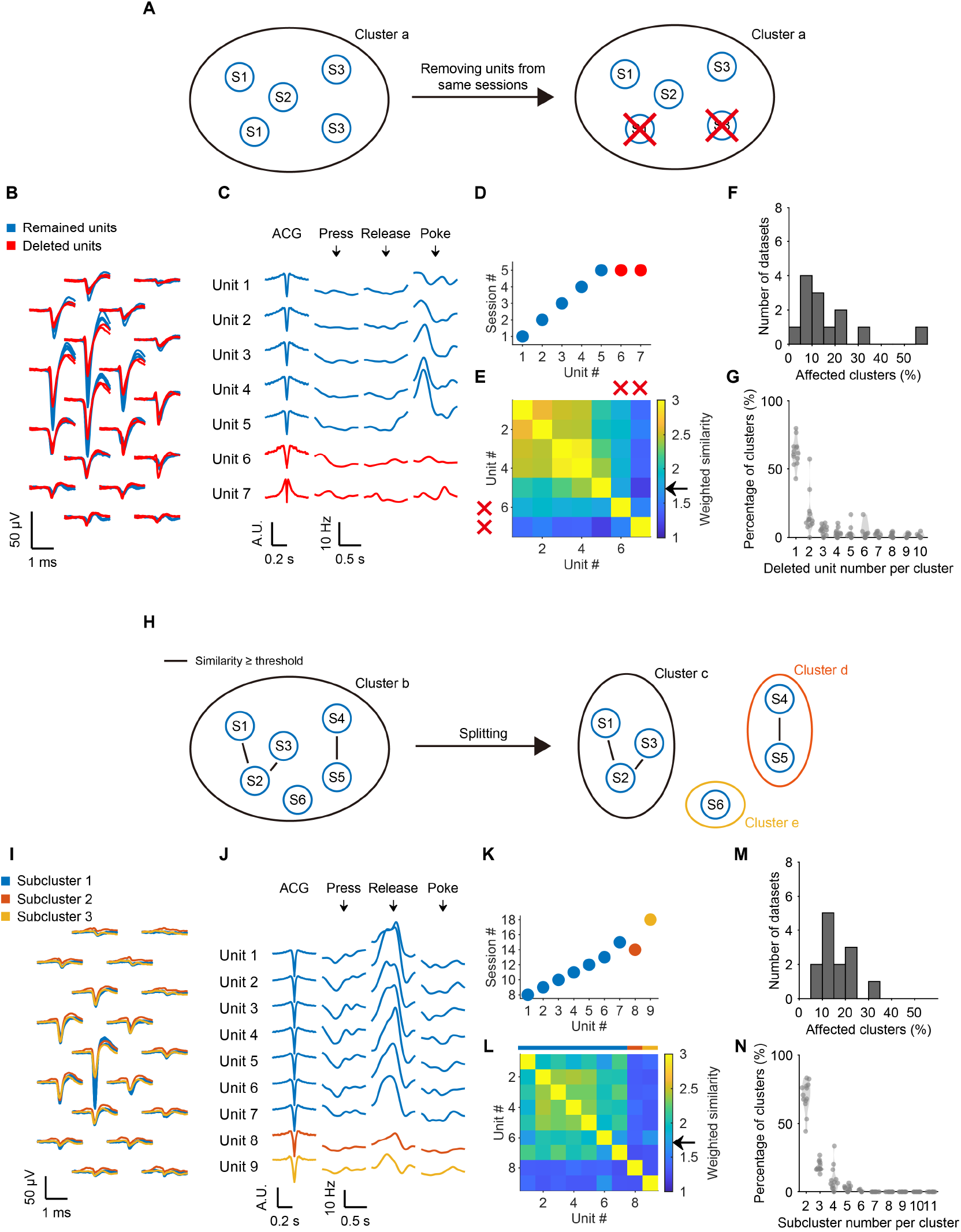
Autocuration for quality control of HDBSCAN clustering results. (A) Schematic of Step 1, in which each cluster is constrained to one unit per recording session. When more than one unit from the same session appears, the units with the lowest average similarity to the other units are removed. (B–E) Example cluster containing multiple units from the same session. (B) Overlaid waveforms, with removed units highlighted in red. (C) Smoothed autocorrelograms and PETHs for all units in the cluster. (D) Session origins for each unit. (E) Pairwise similarity matrix; the arrow on the colorbar indicates the threshold derived from LDA. (F) Distribution across datasets of the proportion of clusters affected in Step 1. (G) Distribution across datasets of the number of units deleted per cluster in Step 1. (H) Schematic of Step 2, in which clusters are split into multiple subclusters if subsets of units exhibit higher mutual similarity than to the remaining units, based on the LDA-derived threshold. (I–L) An example cluster that is split into three subclusters in Step 2; panels follow the same layout and notation as in (B–E). (M) Distribution across datasets of the proportion of clusters affected in Step 2. (N) Distribution across datasets of the number of subclusters per original cluster in Step 2.

**Figure S4.**
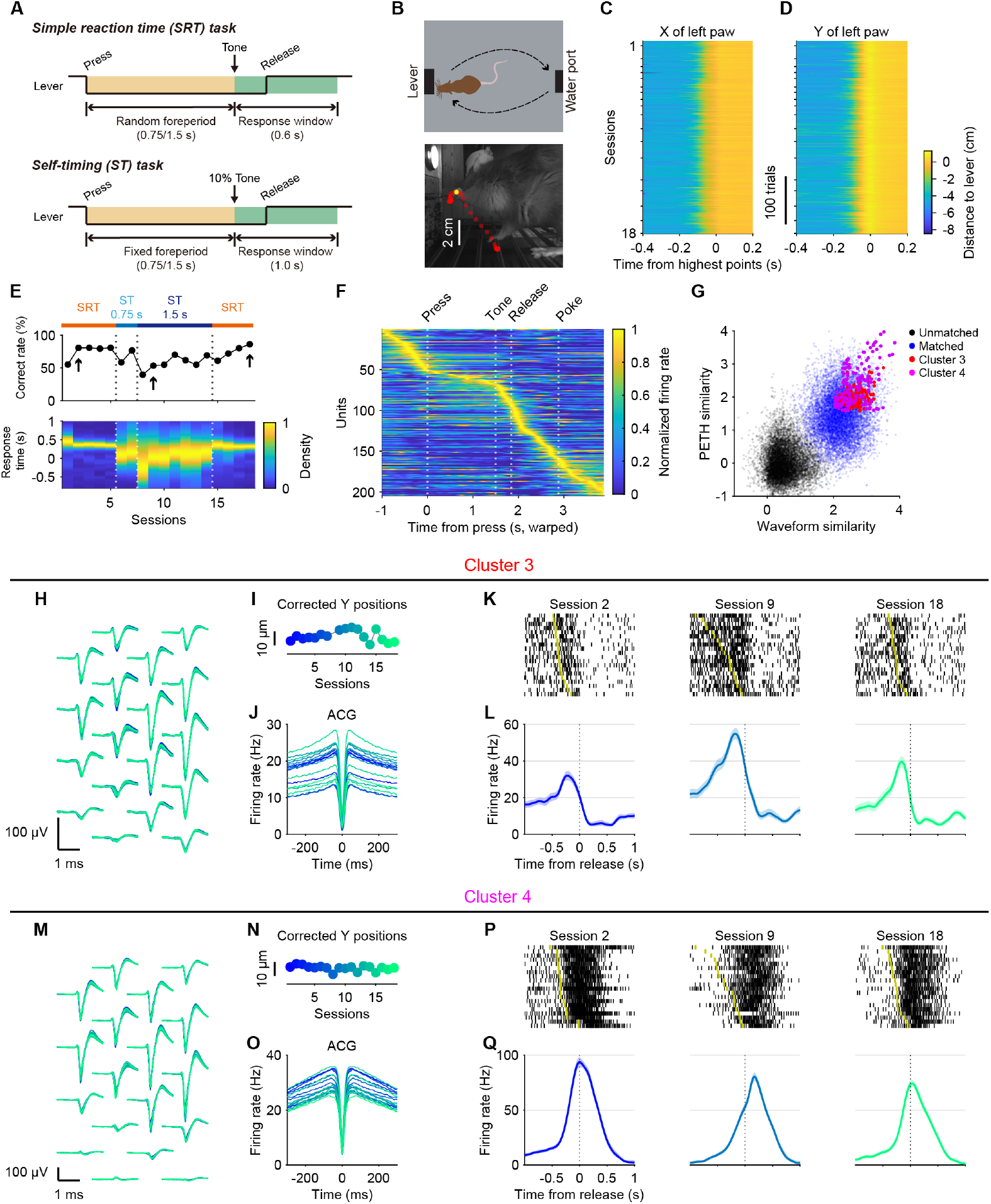
Behavioral paradigm, performance, and example neurons recorded across sessions in an M2 recording. (A) In the simple reaction time (SRT) task, the rat pressed and held a lever for a randomly selected foreperiod (0.75 s or 1.5 s, chosen on each trial) before releasing it within a 0.6-s response window following a brief tone (0.25 s). In the self-timing (ST) task, the rat released the lever after a fixed foreperiod (0.75 s or 1.5 s, constant within a session) within a 1-s response window, with a tone presented in only 10% of trials. (B) Top panel: schematic of the behavioral box in top view. Bottom panel: trajectory of the rat’s forepaw during a lever press, with red points indicating left paw positions and a yellow point marking the peak position. The interval between successive points is 10 ms. (C, D) Left paw tracking across trials and sessions, with color denoting horizontal (X) or vertical (Y) position relative to the lever. (E) Training protocol and behavioral performance across 18 sessions. Top panel shows the sequence of training phases, with the number beneath “ST” indicating the fixed foreperiod used in the ST task. Middle panel plots the proportion of correct trials across sessions. Bottom panel displays the distribution of response times, defined as the duration from the end of the foreperiod to lever release (for premature releases, this value is negative). (F) Time-warped population PETHs for 1.5-s foreperiod trials in a single session (session 2 in the SRT task). PETHs were aligned and linearly warped between lever press, lever release, and nose poke. Units were ordered by the positions of their peak modulation. (G) PETH similarity versus waveform similarity for all pairs of units across 18 sessions, with red and purple dots highlighting unit pairs from the example neurons (153 pairwise comparisons per neuron). (H–J) Same layout as in Figure 4F–H for a different neuron (Cluster 3). Spiking activity is aligned to the time of lever release. (K) Spike raster for 1.5-s foreperiod trials aligned to the lever release. Yellow ticks indicate the end of the foreperiod. (L) PETHs aligned to the time of lever release. Shaded areas indicate the 95% confidence intervals. (M–Q) Same layout as (H–L) for a different neuron (Cluster 4).

**Figure S5.**
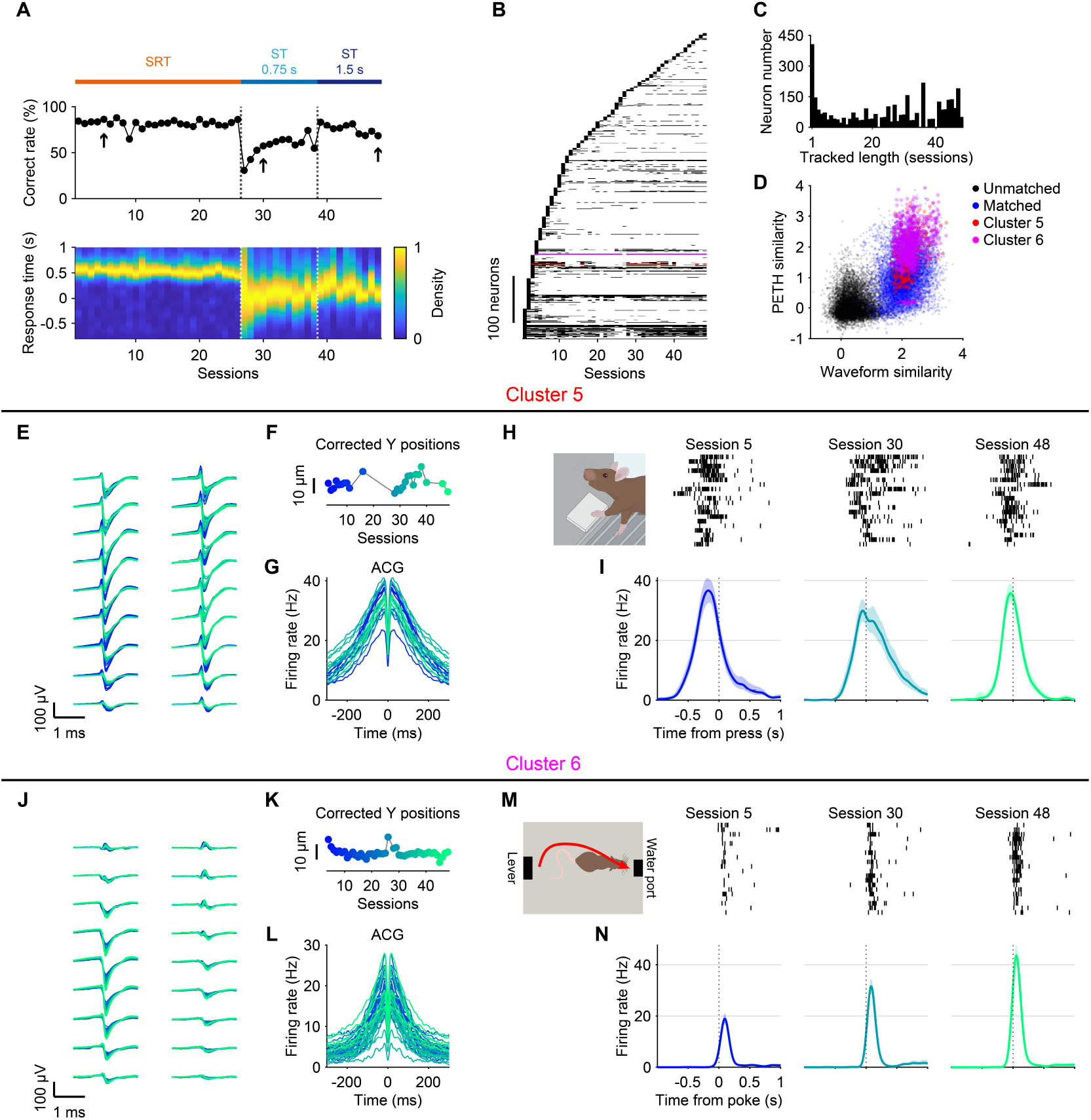
Example neurons recorded across sessions in a DLS recording. (A) Same layout as in Figure S4E for a different animal. (B) Presence of unique neurons across sessions, ordered by first appearance. Red and purple lines indicate the example neurons. (C) Distribution of tracked length (number of sessions in which each neuron was recorded) for all neurons. (D) PETH similarity versus waveform similarity for all pairs of units across 48 sessions, with red and purple dots highlighting unit pairs from the example neurons. (E–I) Same layout as Figure S4H–L for a different neuron (Cluster 5), with spike rasters and PETHs aligned to the time of lever press (J–N) Same layout as (E–I) for a different neuron (Cluster 6), with spike rasters and PETHs aligned to the time of nose poke.

**Figure S6.**
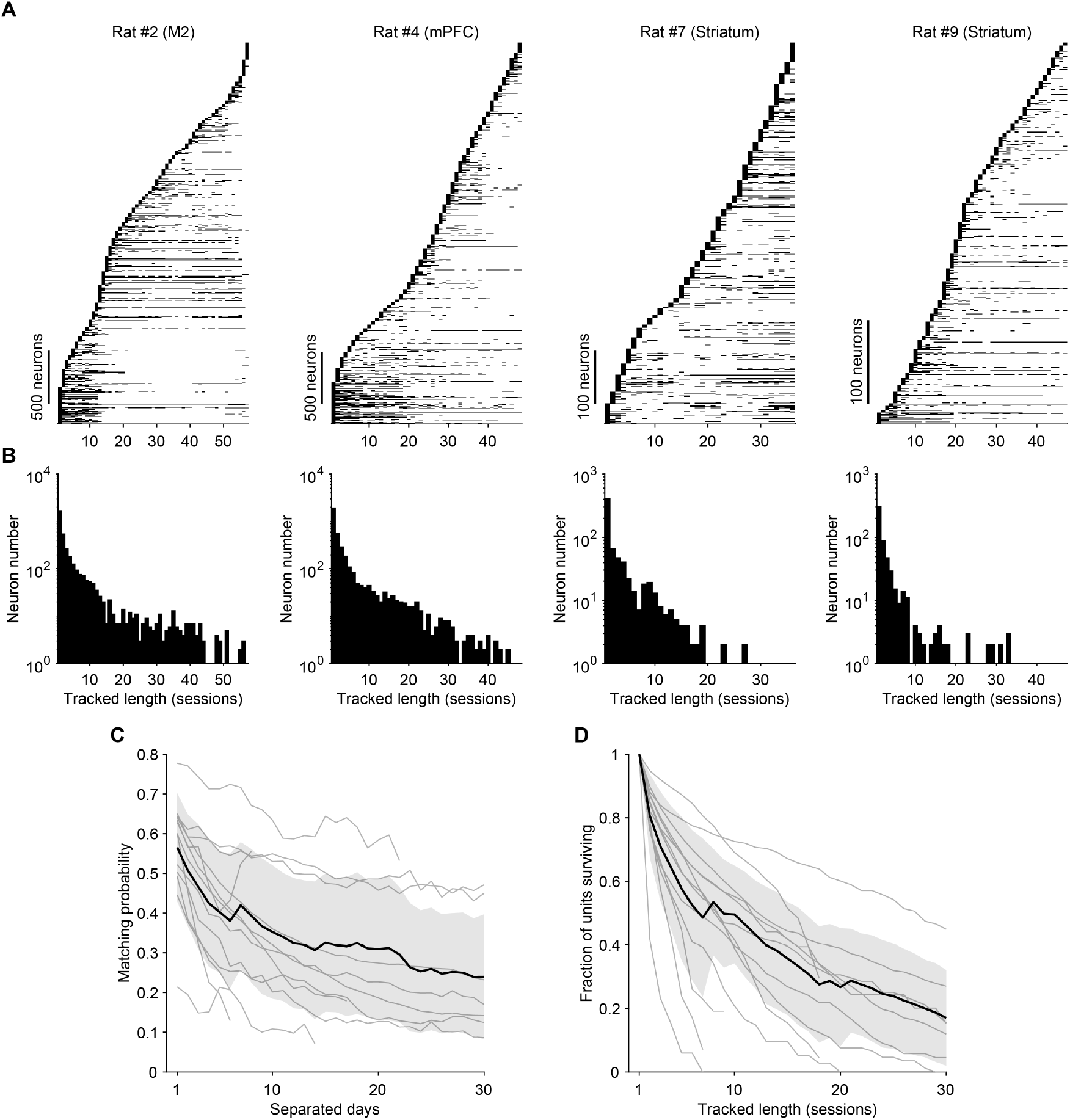
Tracking performance across animals. (A) Presence of unique neurons across sessions detected by DANT in four additional animals. (B) Distribution of tracked length for detected neurons. (C) Matching probability of identifying the same unit between two recordings separated by *n* days across all datasets. Black lines represent mean values, and shaded areas represent standard deviations. (D) Proportion of units that could be tracked for at least *n* sessions across all datasets. Black lines represent mean values, and shaded areas represent standard deviations.

**Figure S7.**
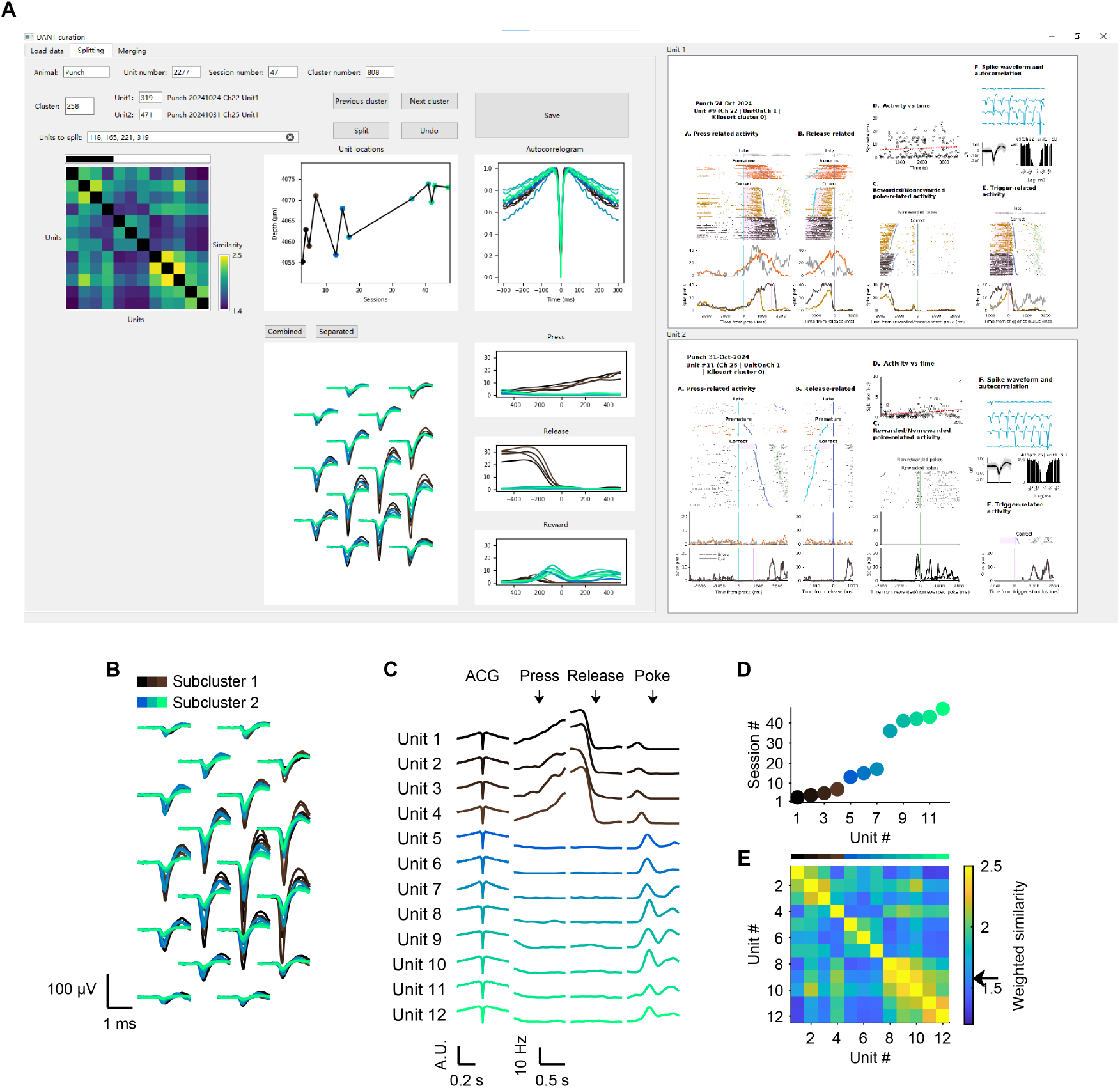
Manual curation of DANT results: an example cluster split into two subclusters with different functional properties. (A) Graphical user interface (GUI) used for manual curation of DANT results. The GUI allows users to visualize and curate the clusters identified by DANT. Displayed features include a similarity matrix between units, unit locations across sessions, motion-corrected waveforms, autocorrelograms, PETHs for three task events, and custom figures illustrating each unit’s functional tuning. The example cluster from Rat #9 was manually divided into two subclusters. Subcluster 1 (the first four units displayed in black) exhibited ramping activity after lever press and before lever release, in clear contrast to subcluster 2 (units displayed in a blue-to-green colormap), which was primarily activated prior to nose poke for reward. The two custom figures on the right correspond to the last unit of subcluster 1 and the first unit of subcluster 2. (B–E) The example cluster from panel (A) is shown using the same layout as in Figure S3B–E.

**Figure S8.**
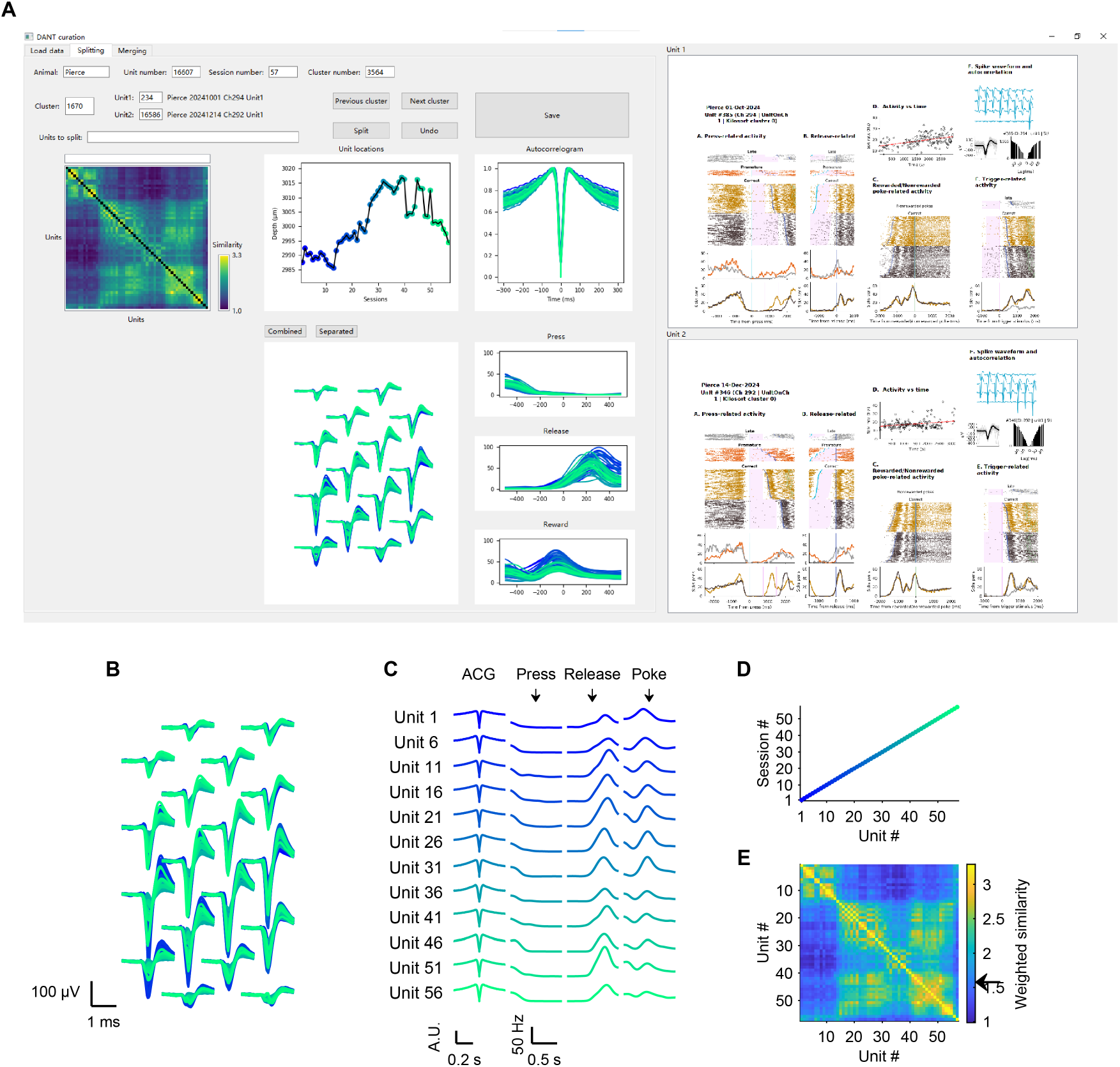
Manual curation of DANT results: an example cluster with consistent functional properties. Same layout as in Figure S7, but showing a cluster from Rat #2 across 57 sessions in which all units passed manual inspection, and no units were excluded.

**Figure S9.**
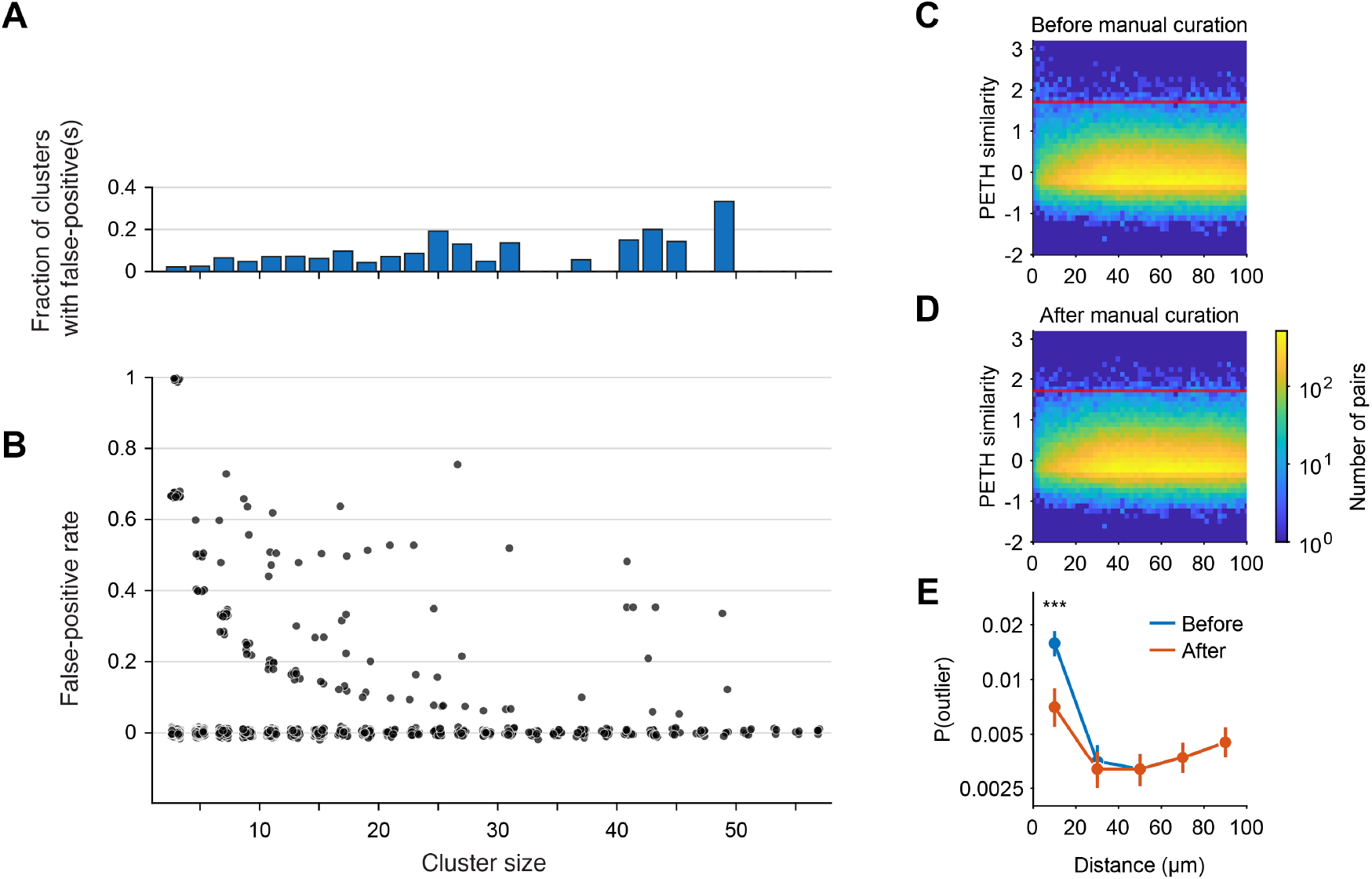
False positives and false negatives identified by manual inspection. (A) Fraction (in each bin) of clusters containing at least one false positive, plotted against cluster size. (B) False-positive rate for each inspected cluster, plotted against cluster size across all datasets. (C) Heatmap of PETH similarity for unmatched unit pairs identified by DANT across consecutive days, plotted as a function of inter-unit distance. Four datasets from motor cortex and dorsolateral striatum recordings were included. Pixel intensity reflects pair counts. The red line represents the threshold used to identify outliers. (D) Same as (C), but after manual curation, including merging clusters that were presumed to be the same neurons. (E) Proportion of outliers among unmatched pairs before and after manual curation, grouped into 20-μm distance bins. Error bars indicate 95% confidence intervals. ***: *p* < 0.001.

**Figure S10.**
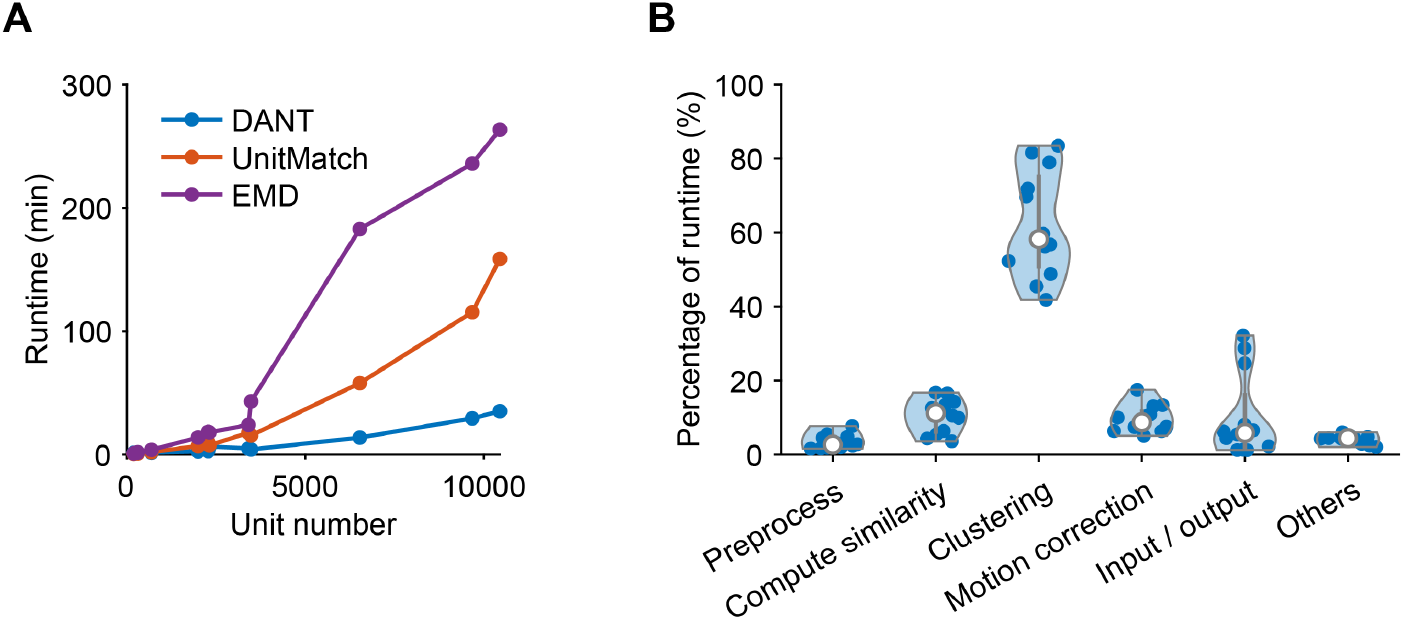
Evaluating runtime performance. (A) Total runtime across 11 datasets for different methods. (B) Breakdown of runtime proportions for individual processes within DANT.

**Figure S11.**
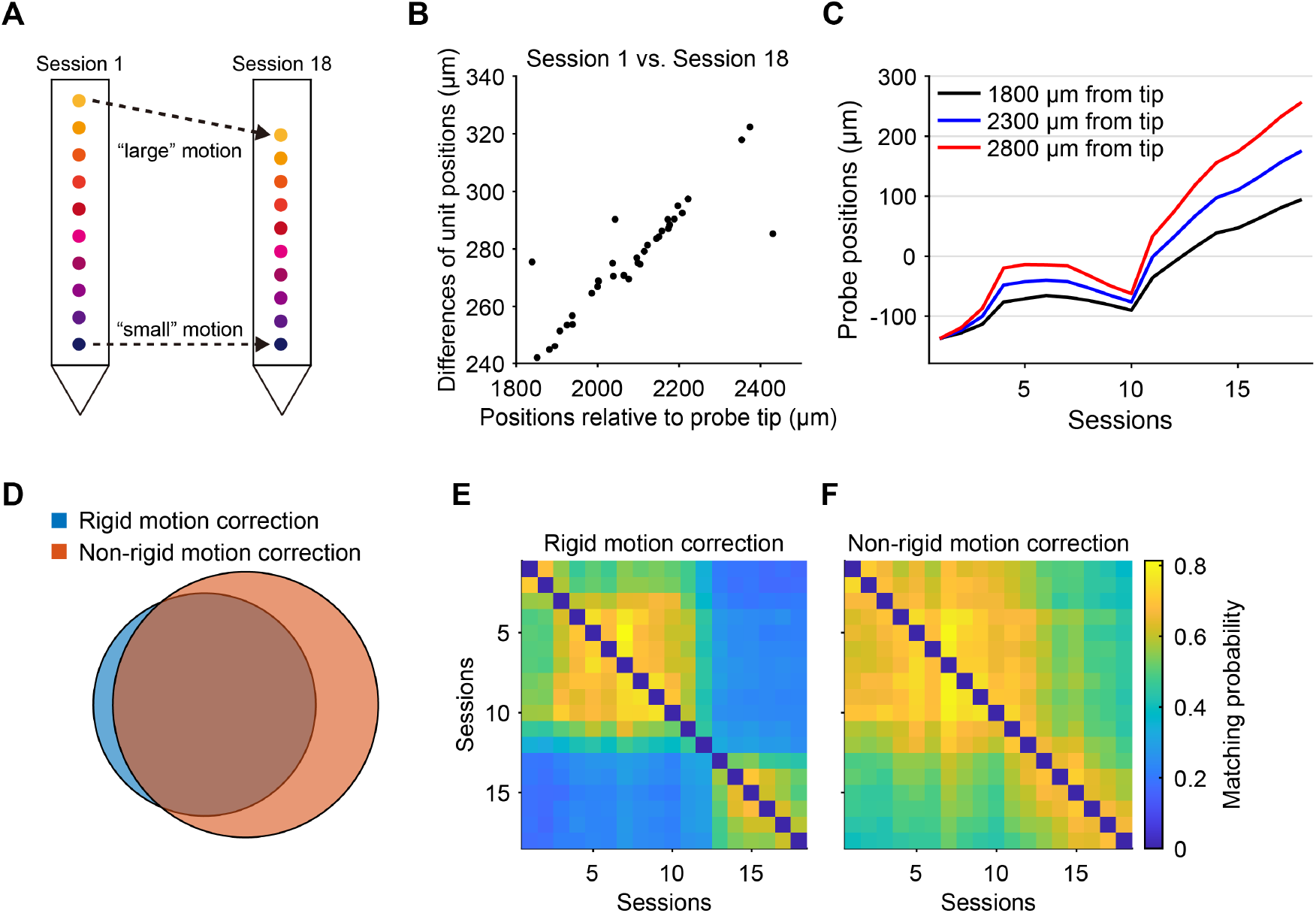
Non-rigid motion correction. (A) Schematic illustration of depth-dependent non-rigid motion. (B) Motion between session 1 and session 18 for matched unit pairs at different positions along the probe in Rat #3. The x-axis represents the average positions of same-neuron pairs. (C) Probe motion estimates obtained by DANT. (D) Venn diagram showing the overlap of matched units identified by DANT with rigid versus non-rigid motion correction. (E–F) Matching probability matrices showing tracking results of DANT under rigid and non-rigid motion correction.

Rigid motion correction is often insufficient to capture the full extent of probe displacement. Several datasets exhibit pronounced depth-dependent motion along the probe (Figure S11A,B). For example, in Rat #3, matched pairs identified by DANT under rigid correction show a strong linear correlation between unit positions and their relative displacements across sessions (Pearson’s *r*=0.999; Figure S11B). The displacement difference between the top and bottom regions exceeds 160 μm (Figure S11C), making accurate tracking difficult because of imprecise waveform correction. Non-rigid correction is thus required. Conventional approaches address this issue by dividing the probe into “blocks” and correcting motion within each block. However, this strategy is ill-suited for neuron tracking because non-uniform unit distributions along the probe often leave some blocks with insufficient data for reliable motion estimation.

To address these limitations, we developed a modified algorithm for non-rigid motion correction. Following the definition used in rigid motion estimation, we additionally define the depth *d*^*i*^ for the *i*-th matched unit pair as

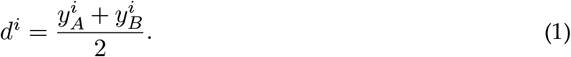

where 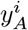 and 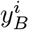 denote the depths of the matched units in sessions 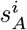 and 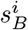, respectively. We then introduce two vectors, *k* and *b*, each of length *N*_*s*_ (the number of sessions), representing depth-dependent motion (slope) and overall probe displacement (offset) across sessions. These parameters are estimated by solving

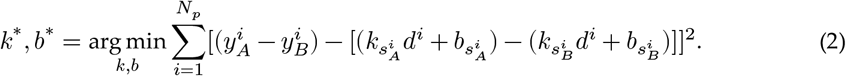

This formulation reduces to rigid correction when all *k* values are set to zero. To ensure proper minimization, *k*_1_ is fixed at zero, assuming no non-rigid motion in the first session. The solution *b*^∗^ is then mean-subtracted via

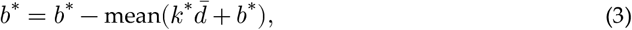

where

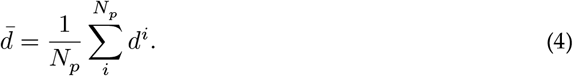

Instead of a direct readout of probe positions as in the rigid case, the probe position *p*^*i*^(*d*) at the depth *d* in session *i* is computed as

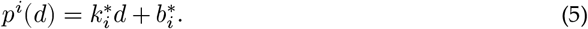

The subsequent waveform correction step proceeds as in the rigid case. Applied to Rat #3, this method revealed greater magnitude of motion in regions distal to the probe tip (Figure S11C). Compared with rigid motion correction, the non-rigid algorithm identified 42.2% more matched unit pairs (rigid: 23121 matches, with each neuron tracked for 8.1 ± 5.2 sessions; non-rigid: 32895 matches, with each neuron tracked for 11.1 ± 5.5 sessions, mean ± SD; Figure S11D–F). The matching probability from session 1 to session 18 increased from 0.17 to 0.38 (Figure S11E,F). Together, these results demonstrate that non-rigid motion correction algorithm substantially improves tracking performance in datasets with depth-dependent probe motion.

